# Delayed cerebrovascular dysfunction and social deficits after traumatic brain injury

**DOI:** 10.1101/2024.02.15.580515

**Authors:** Aditya Singh, Steven Gong, Anh Vu, Scott Li, Andre Obenaus

## Abstract

Traumatic brain injury (TBI) survivors face debilitating long-term psychosocial consequences, including social isolation and depression. Acute TBI modifies neurovascular physiology and behavior but a gap in our understanding are the chronic physiological implications of altered brain perfusion on behavioral activities, particularly social interactions.

We investigated longitudinal functional vascular networks across the brain for 2-months post-TBI and its impact on social behavior. Adult C57/BL6 male mice received a moderate cortical TBI. Behavior (foot-fault, open-field, 3-chamber social preference) was assessed at baseline, 3-, 7-, 14-, 30-, and 60-days post injury (dpi) followed by magnetic resonance imaging (MRI, 9.4T). Anatomical MRI (T2-weighted), dynamic susceptibility contrast (DSC) perfusion weighted MRI (PWI) were acquired at each temporal epoch. After the final 60dpi MRI, animals underwent transcardial perfusion fixation to map angioarchitecture. MRI data were analyzed using standardized protocols followed by cross-correlations between social behavior, cerebral perfusion, and vascular metrics.

Social behavior deficits at 60dpi emerged as reduced interaction with a familiar cage-mate (partner). We observed multiphasic decrements in cerebral blood flow (CBF) encompassing lesion and perilesional cortex where acute reductions at 3-14dpi partially recovered by 30dpi, followed by significant reductions in perfusion at 60dpi. The CBF perturbations extended antero-posteriorly from the ipsilateral TBI impact site but also adulterated contralateral brain regions. CBF reductions impacted regions known to regulate social behavior including hippocampus, hypothalamus, and rhinal cortex. Alongside perfusion deficits at 60dpi, social isolation in TBI-mice emerged with a significant decline in preference to spend time with a cage mate. Cortical vascular density was also reduced corroborating the decline in brain perfusion and social interaction.

Thus, the novel temporal neurovascular loss, and subsequent recovery followed by chronic decrements are broadly reflected by social interaction perturbations. Our correlations strongly implicate a linkage between vascular density, cerebral perfusion, and social interactions, where early evaluation can potentially predict long-term outcomes. Thus, our study provides a clinically relevant timeline of alterations in functional vascular recovery that can guide research for future therapeutics.

## Introduction

Traumatic brain injury (TBI) afflicts an estimated 70 million people across the world annually with both age and sex as important determinants of severity and outcomes.^1, 2^ Mild to moderate TBI are the most common type of injuries among civilians, professional sports personnel, and military service members.^3, 4^ Behavioral, physiological and psychosocial deficits persist for extended periods after the initial injury and include depression, post-traumatic stress, sleep, Alzheimer’s disease, and dementia related disorders amongst others.^5–8^ Despite advances in our understanding of the pathophysiological processes underlying TBI, there is a gap in the linkage between behavioral outcomes and cerebrovascular alterations following TBI.

Social interactions are impaired after TBI with reductions in interpersonal communication,^9^ time with friends and families,^10, 11^ and is evident in children^12, 13^ and adults.^14, 15^ Social behavior is also diminished in adult mice after a single^16, 17^ or repeated-mild TBI,^18, 19^ and in pediatric models of TBI.^20^ Social isolation in rodents reflects increased neuropathology after TBI,^21^ however, increased social interaction post-injury is known to facilitate recovery.^22^ Lacking are studies that examine how TBI modifies sociability and its relationship to disrupted cerebrovascular morphology and function.

We and others have reported acute vascular alterations following TBI spanning chronically reduced microvasculature,^23–25^ and global reductions in cerebrovascular reactivity and tone.^26–28^ TBI induced changes in the vasculature are associated with motor and cognitive behavioral deficits,^29, 30^ while others have reported no change at long-term after injury.^31^ Autism spectrum (ASD) subjects, which manifest deficits in social interactions, exhibited CBF reductions that correlated with the severity of behavioral alterations.^32^ Further, intranasal treatment with oxytocin increased blood flow across social processing brain regions^33^ and monitoring vascular metrics has been proposed as a biomarker for social interactions.^34^

To address the paucity of knowledge linking physiological CBF, underlying angioarchitecture, and social behavior deficits after moderate TBI, we undertook a longitudinal study in adult male mice. Specifically, we tested the hypothesis that long-term cerebrovascular deficits facilitate social dysfunction. We report that temporal vascular flow and morphology initially recover but ultimately decline by 2 months post TBI which are mirrored by social interaction deficits . Our novel study provides the basis for future preclinical and clinical interventional studies targeting social psychopathologies following acquired moderate TBI.

## Materials and methods

### Animals

All experiments were conducted using ARRIVE guidelines and animal use was approved by the University of California Irvine Animal Care and Use Committee. Adult C57/B6 male mice (JAX#000664, 2-3months old) from Jackson Laboratory were group housed with 12hour light/dark cycle in ventilated cages and acclimated for a minimum of 7d after arrival. Male C57BL/6 mice underwent either sham surgery (n=11) or a moderate TBI (n=9) followed by longitudinal behavior and perfusion MRI across a 60d post injury (dpi) time course (Fig. 1A, B). A subset of sham and TBI mice (n=6/group) were relegated for behavior. Two of nine TBI mice died at 30dpi. Sham and TBI mice maintained similar weight profiles (Supplementary Fig. 1).

**Figure 1.**
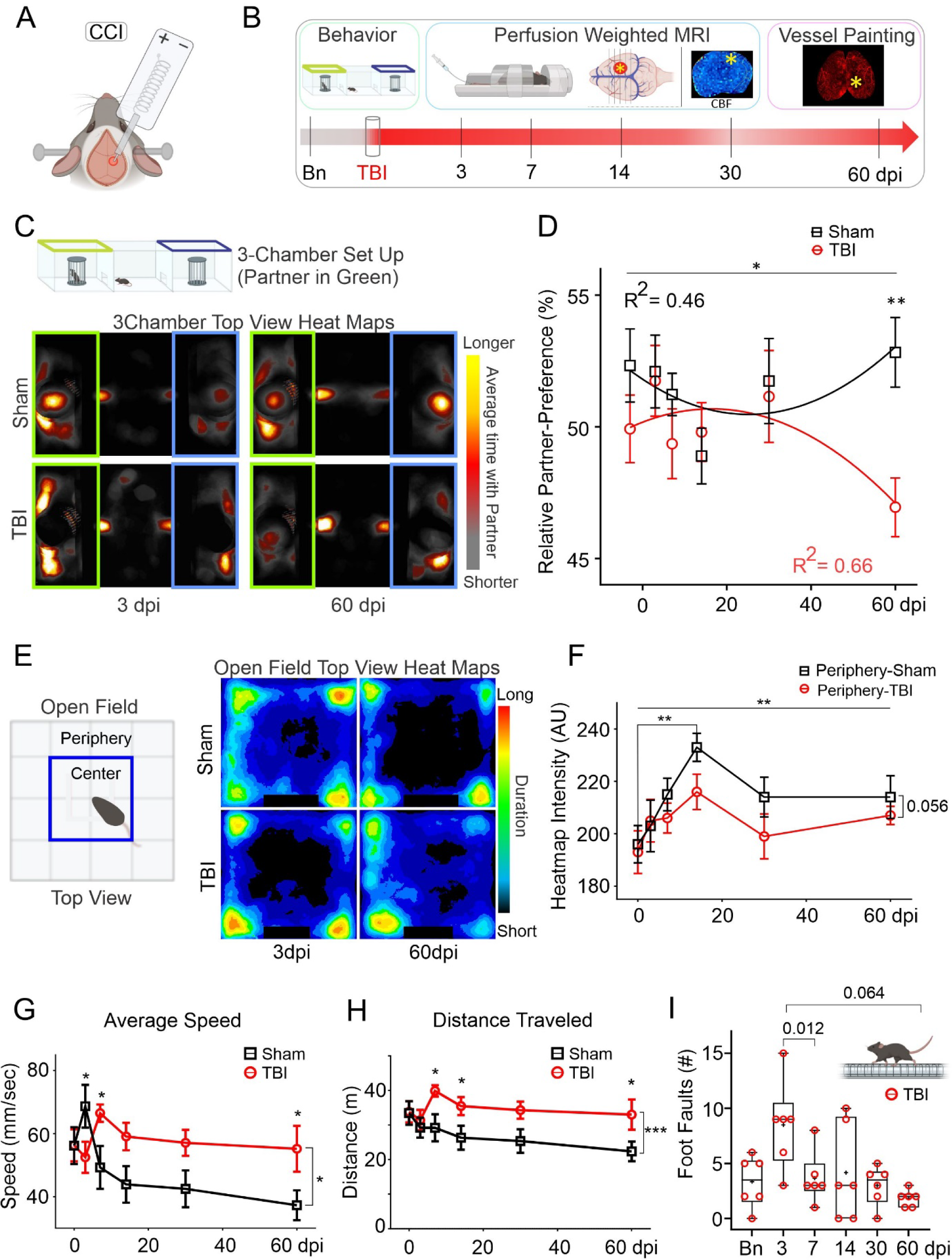
Chronic social and exploratory behavior changes following TBI. (A) Cortical contusion injury (CCI) in adult male mice centered at somatosensory and motor cortices. **(B)** Behavioral and MRI experimental timeline with vessel painting at 60 days post injury (dpi). **(C)** Heat maps of 3-chamber social behavior utilized a known cage-mate mouse (partner) and illustrate increased cumulative time spent by sham (top) and TBI mice (bottom row) at 3 and 60dpi **(D)** Relative partner-preference (RPP) is significantly reduced at 60dpi in TBI (red circles) relative to sham mice (black squares) (2wANOVA - Injury factor - *F*(1,60)=5.02, \**p*=0.029, Tukey’s post-hoc 60dpi. sham vs TBI - ** *p*=0.002). **(E)** Open-field arena schematic (left) with center and periphery zones. Heatmap of average time spent at 3- and 60dpi for sham (top row) and TBI animals (bottom row) illustrates increased center time after TBI. **(F)** Sham mice (black, squares) spent significantly more time in periphery at 14dpi vs. baseline (2wANOVA: *F*(5,61)=0.006, Tukey’s post hoc \*\**p*=0.008) but reduced time in TBI mice (Injury factor, 2wANOVA, *F*(1, 61)=3.80, *p*=0.056). **(G)** TBI mice in open field exhibited increased average speed compared to shams (2wANOVA, injury factor, *F*(1, 61)=6.44, \**p*=0.014). **(H)** Total distance travelled in open field was also significantly increased in TBI mice vs shams (2wANOVA, injury factor, *F*(1, 61)=14.4, \*\*\**p*=0.0003) at 7, 14 and 60dpi (*p<0.05). **(I)** Sensorimotor tests in TBI mice at 3dpi exhibited increased foot-faults with modest longitudinal recovery. (dpi - days post injury, CCI - Cortical contusion injury, TBI - Traumatic brain injury, partner - cage mate mouse, CBF - cerebral blood flow, Bn - baseline, bright asterisk on coronal and axial view of the brain - TBI impact site.)

### Traumatic brain injury (TBI)

TBI with controlled cortical impact (CCI, Fig. 1A) was induced as previously reported^25, 35^ and detailed in the supplementary materials. Briefly, anesthetized (isoflurane 1-3%) mice were maintained at 37°C and then placed in a stereotaxic device. Under aseptic conditions, a scalp incision was made, underlying connective tissue retracted and a 5mm craniotomy (bregma AP –1.25cm, ML +1.25cm) was performed to expose the brain. A 1.5mm impactor tip was zeroed to pial surface and electromagnetic impactor was discharged (Leica, NeuroscienceTools, O’Fallon) with the following parameters: 1mm depth, 200ms dwell-time, speed 5m/s. Extravascular bleeding was immediately wicked away and the skin was sutured closed without replacing the bone. Buprenorphine (100ng/g body weight, intramuscular) was injected, and mice were returned to a warmed chamber until ambulatory. Sham mice underwent anesthesia exposure for the same duration.

### Behavioral Paradigms

Mice were carefully handled to habituate with experimenter 1-2 days before testing.^36^ Behavioral testing sequentially included foot-fault (FF), open-field (OF), 3-chamber social preference (3Ch) tests prior to each magnetic resonance imaging (MRI) session for a subgroup of animals (*n* = 12 for baseline, 3, 7, 14 and 30dpi, *n* = 16 at 60dpi). Mice in their home cages were acclimatized to the behavior room for 5-10 min before onset of testing. Mice were allowed to rest for 10min in their home cage between testing paradigms. All apparatus were disinfected before and after each mouse. Extended behavioral details are delineated in the Supplementary materials.

### Foot-fault (FF)

Mice were individually placed at one end of a grid (47.5 x 29.5cm, 25 beams, 1.5cm apart) and allowed to walk freely until they reached home cage at the other end of the grid or 120seconds, whichever occurred first. Foot slips through the grid were manually counted by two blinded experimenters.

### Open Field (OF)

Mice were placed in the center of an open field arena (30 x 30cm^2^) and allowed free exploration for 10min (top view webcam recording).

### 3-Chamber Social Preference (3Ch)

Our 3Ch test utilized a known cage-mate (partner) being placed inside a wireframe enclosure in one of the peripheral chambers and alternated between each session^37^. The peripheral chambers were connected to the central chamber (15.5 x 28.50cm^2^) with manual sliding doors. The test mouse was placed in the central chamber with closed doors for 5min and then doors were opened to allow free access to both peripheral chambers for 10min. The behavior was video recorded for offline analysis.

### Behavior Analyses

All semi-automated image analyses or manual scoring were blinded to injury condition. We utilized multiple software, including Fiji^38^ for OF and 3Ch, and AnimalTA^39^ for OF animal tracking to estimate speed and distance. Videos were cropped to isolate the identical regions of interest (ROIs) for OF and 3Ch (Supplementary Fig. 2). FIJI’s image adjust algorithm was used with automated minimum threshold for animal (red) and background (dark) detection, to generate binary masks. Thresholded stacks were averaged as heat-maps (Fiji), with pixel-intensity representing time^40^ allowing measurement of time spent in partner vs no-partner chambers, termed relative partner-preference (RPP). Manual scored behavior utilized BORIS^41^ to derive, a) absolute partner-interaction time (API) defined as total time the test mouse was facing the cylindrical enclosure with the partner mouse, and b) relative partner-interaction time (RPI), defined as the ratio of time spent interacting with (pointed towards) partner enclosure vs. total interaction time across both partner and non-partner enclosures.

### MRI

In vivo longitudinal MRI was performed (Fig. 1B) at baseline and after TBI induction (3, 7, 14, 30, 60dpi) on a horizontal 30cm bore, 9.4Tesla MR scanner (Bruker Avance) equipped with a 72mm diameter volume excitation RF coil. Our perfusion weighted imaging (PWI) MRI methods are published^42^ and described in more detail in the supplementary materials. Succinctly, mice were anesthetized (2% isoflurane) and tail veins were cannulated to facilitate contrast injection (0.1mmol/kg Gadoterate Meglumine diluted with sterile saline, Dotarem, Guerbet, Princeton, NJ). Mice were then placed in MRI and the following sequences were acquired: T2-weighted (T2), T1-weighted images (T1), PWI during which Gd was infused (1ul/g body weight), and susceptibility-weighted MRI (Supplementary Table 1 for MRI sequence details).

### MRI Image Analysis

Detailed MRI processing methods are reported in Supplementary materials. PWI MRI was processed using Jim software (V9.1, Xinapse Systems Ltd, Essex, UK) using the Brain Perfusion tool to automatically derive the arterial input function (AIF) curves which were manually reviewed for typical AIF profiles. AIF curves from each sham mouse across all six time points were averaged for a group average AIF^43^ and used to calculate cerebral blood flow (CBF, ml/100g-tissue/min) and cerebral blood volume (CBV, %tissue).^44^ TBI animals used individual AIF curves to calculate CBF and CBV to account for variability due to injury. In Jim software an in-house mouse atlas was applied to CBF and CBV parametric maps, values were extracted and summarized in Excel.

### Vessel Painting and Analyses

To visualize cortical angioarchitecture we utilized our vessel painting protocol as previously reported (see supplementary materials).^35^ Briefly, 1,1’-Dioctadecyl-3,3,3’,3’- Tetramethylindocarbocyanine Perchlorate (DiI, D282, Invitrogen, Carlsbad) was delivered via intracardiac injection prior to,) infusion was performed prior to transcardial fixative perfusion at 60dpi and brains were extracted after 24hr post fixation, rinsed in 4% PFA for 24hours and stored in PBS+0.02% sodium azide until microscopic acquisition. All animals had successful staining of the cortical vasculature (n=8 TBI, n=11 sham).

Vascular image acquisition and analysis protocols have been previously published.^23, 35^ Briefly, the bilateral axial cortices were imaged at 2X magnification using an epifluorescent wide-field microscopy (BZ-X810, Keyence Corp., Osaka) and 10x magnification images of the middle cerebral artery (MCA) of the ipsilateral hemisphere. Classical vascular analysis for vessel density, junctional branch points, total end points, and average and total vessel length were obtained using the Fiji plugin, Angiotool.^45^ Analysis focused on lesion and perilesional regions (see Fig. 4A). Fractal analyses for vascular complexity was also performed using Fiji Fraclac plugin to obtain local fractal dimensions (LFD).

### Statistics

Behavioral indices, MRI, and vessel painting derived values were imported into MS-Excel. MRI data were filtered for outliers using interquartile ranges. Statistical testing, Pearson’s correlation coefficient estimations, and plotting were performed using MS-Excel or Prism 9.0 (GraphPad, San Diego). Scatter plots used box and whisker graphs with mean and error bars (minimum to maximum values) where the box bounding represents 25th and 75th percentiles. In line graphs error bars are plotted as standard error of mean (SEM). Two-way ANOVA (2wANOVA) with Tukey’s Post hoc test was used for statistical comparisons, unless specified otherwise. Statistical significance was noted at \**p*<0.05, \*\**p*<0.01, or \*\*\**p*<0.001, with trending as *p*<0.1.

## Results

### Long-term reduction in social behavior after TBI

Social behavior using a 3Ch paradigm with cage-mate mice illustrate similar relative partner-preference (RPP) at 3dpi but dramatic reductions at 60dpi (Fig. 1C, Supplementary Fig. 3A- D). Semi-automated quantitative image analyses confirmed a significant RPP decrease in cage- mate interaction (2wANOVA, Injury: *F*(1,60)=5.02, \**p*=0.029) with significant post-hoc reductions in TBI animals at 60dpi (*Adj. **p*=0.002) (Fig. 1D, Supplementary Fig. 4). Manual video scoring confirmed similar RPP profiles (Supplementary Fig. 3A-D). The social deficits were independent of motor deficits, as evident by FF and OF tests, confirming comparable speeds with higher exploratory drive likely reflecting increased risk-taking behavior in TBI-mice vs. shams (Fig. 1E-H).

### Increased exploratory behavior after TBI

Increased risk-tasking in TBI mice was evident with significantly reduced OF periphery activity as early as 14dpi through to 60dpi relative to shams (Fig. 1E, F) (2wANOVA, Injury, *F*(1,61)=3.80, *p*=0.056; Timepoints, *F*(5,61)=3.64, \*\**p=*0.006). Post-hoc, sham mice spent more time in periphery at 14dpi compared to baseline (**p=0.008). Ratio of time spent in center/periphery found no significant differences (Supplementary Fig. 3F).

TBI mice exhibited higher speeds and distance travelled from 7dpi onwards across all the timepoints with significant ‘time X injury’ interactions for speed (Fig. 1G-H, Supplementary Fig. 3E-H); (Fig. 1G, average speed, 2wANOVA, Injury, *F*(1,61)=6.44, \**p*=0.014; Timepoint, *F*(5,61)=1.91, *p*=0.1, Interaction *F*(5,61)=2.99, \**p*=0.018); (Fig 1H, total distance, 2wANOVA, Injury, *F*(1,61)=14.4, \*\*\**p*=0.0003; Timepoint, *F*(5,61)=1.28, *p*=0.28, Interaction *F*(5,61)=1.01, *p*=0.42). TBI animals had significantly reduced speed at 3dpi (*Adj.*p*=0.045) but elevated at 7dpi (*Adj. *p*=0.027) and 60dpi (*Adj.*p* =0.027) vs. sham mice. Distance travelled was elevated at 7dpi (*Adj.*p*=0.017), 14dpi (*Adj. *p*=0.048), and 60dpi (*Adj.*p*=0.022) for TBI vs. sham mice.

### Early sensorimotor deficits post-TBI recover with time

Sensorimotor failures assessed using FF testing were increased in TBI compared to sham mice at 3dpi (Supplementary Fig. 5, 1wANOVA, *F*=3.99, p<0.001, *Post hoc* 3dpi TBI vs. Sham, \*\**p*=0.001), with return to baseline between 7-60dpi (Fig. 1I, rmANOVA, *F*(4.28, 11.8)=4.28, \**p*=0.035, post-hoc: 3- vs 7dpi \**p*=0.012, 3- vs 60dpi *p*=0.064).

### CBF dysfunction mirrors social behavior deficits

Structural T2WI in TBI mice exhibited early edema which resolved over time (3-7dpi) with subsequent cortical thinning at the impact site (14-60dpi; Fig. 2A). Lesion volumes were initially elevated during the edematous phase and then stabilized to ∼10mm^3^ or ∼4% of brain volume (Fig. 2B, C). CBF exhibited regional and global declines that gradually recovered over the initial 30dpi, but then steeply declined at 60dpi (Fig. 2A, D). CBF at the cortical impact site in TBI mice had acute reductions at 3dpi, modest recovery during 7-30dpi, followed by precipitous CBF declines at 60dpi (Fig. 2D, mixed effect 1wANOVA, slice#1: *F*(2.993, 24.55)=3.71, \**p*=0.02, 3- vs 7dpi \**p*=0.036, and 3- vs. 30dpi \**p*=0.046; slice#4: *F*(2.66, 23.38)=5.20, Tukey’s post-hoc: Bn vs 7dpi \**p*=0.031, Bn vs 14dpi ***p<0.001, Bn vs 30dpi \**p*=0.036, and Bn vs. 60dpi \*\**p*=0.006). TBI induced CBF perturbations extended beyond the impact site to adjacent and distant ipsilateral cortical and subcortical regions (Fig. 2A, E) where CBF heatmaps highlight the regional multiphasic nature of physiological recovery. Like the injury site profile (Fig. 2D), distant regions reflected an initial CBF decline at 3dpi, transient recovery at 7-30dpi with a subsequent decline at 60dpi (Fig. 2E). We then examined the relationship between social behavior (RPP) and CBF at 60dpi which demonstrated positive correlations in regions involved in exploratory and social behavior (Fig. 2F-H). Hippocampal CBF was trending positively correlated to RPP (Fig. 2F, *p*=0.08, *R^2^*=0.30) and was significantly correlated in the entorhinal cortex (Fig. 2G, \**p*=0.04, *R^2^*=0.30), but not in somatosensory cortex (Fig. 2H, *p*=0.28, *R^2^*=0.12). Thus, TBI elicits a dynamic profile of tentative recovery followed by regional reductions that correlated to indices of social isolation.

**Figure 2.**
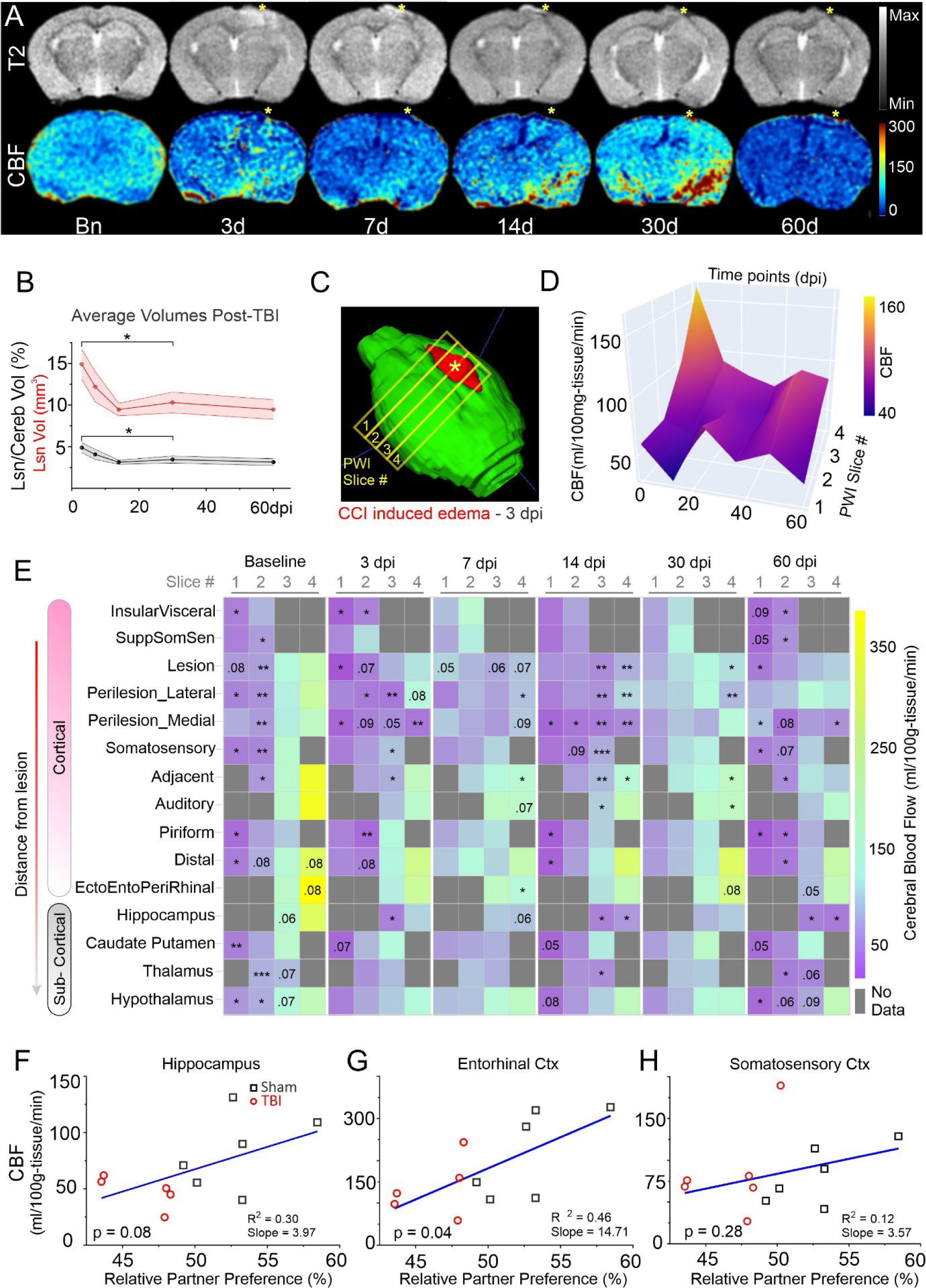
Cerebral blood flow (CBF) recovers but declines at 60dpi. (A) Representative T2-weighted anatomical MR images and CBF maps (ml/100g-tissue/min) from the same TBI mouse illustrates transient decrements, recovery and then followed by precipitous decline at 60dpi. Edema at 3dpi at the impact site (asterisk) resolves and is followed by moderate tissue loss 14-60dpi. **(B)** Lesion volume (mm^3^, red; % brain volume, black) edema increases and stabilizes after edema resolution (1wANOVA – Lesion Volume, *F*(2.17, 18.4)=5.43, *Geisser-Greenhouse’s ɛ*=0.541, \**p*=0.013, *Tukey’s* post-hoc 3- vs. 30dpi \**p*=0.47; Lesion/Cerebrum volume, *F*(2.18, 18.5)=5.37, *Geisser-Greenhouse’s ɛ*=0.544, \**p*=0.013, *Tukey’s* post-hoc 3- vs. 30dpi \**p*<0.05). **(C)** Brain 3D-reconstruction in a TBI mouse (3dpi) illustrates edematous lesion (red). PWI MRI data were collected from four 1mm thick coronal slices. **(D)** Temporal evolution (Basline-60dpi) of CBF at lesion site across antero-posterior slices with acute reductions at 3dpi, recovery followed by declines at 60dpi. **(E)** CBF heatmap depicting longitudinal CBF changes for each slice (columns) with brain regions (rows) sorted by distance from TBI impact site. Statistical significance (t-test TBI vs Sham) is noted (* *p*<0.05, ** *p*<0.01, *** *p*<0.001) as are trending *p*-values. Reduced CBF was evident in anterior slices but increased in posterior slices distant from TBI site. **(F-H)** Correlations between 60dpi CBF and relative partner preference (RPP) in sham and TBI mice in social exploration related brain regions (dorsal hippocampus *p*=0.08, *R*^2^=0.30 **(F)**, entorhinal cortex *p*=0.04, *R*^2^=0.46 **(G)** and somatosensory cortex (*p*=0.28, *R*^2^=0.12, **(H)**).

### Cortical and sub-cortical progression of CBF dynamics post-TBI

We next investigated regional CBF profiles as a function of distance from the impact site (Fig. 3A). The lesion site CBF was significantly lower across the 60dpi epoch, reflecting protracted neurovascular damage after TBI (Fig. 3B, Injury, *F*(1,105)=8.36, \*\**p*=0.005, Timepoints, *F*(5,105)=2.51, \**p*=0.035). After an initial decline, lateral perilesional (somatosensory) cortex CBF progressively increased above shams (Fig. 3C, Injury, *F*(1,106)=4.47, \**p*=0.037, Timepoints, *F*(5,106)=3.77, \*\**p*=0.003). *Post hoc*, CBF at 3dpi was acutely reduced after injury (\*\**p*=0.004). Longitudinally for TBI animals, CBF at 3dpi was lower compared to 30dpi (\**p*=0.026) and 60dpi (\*\**p*=0.003), and 14dpi CBF was lower vs. 60dpi (\**p*=0.042) in TBI animals, demonstrating a multiphasic pattern with long-term increment after initial decline. Notably, the recovery by 30dpi was reflected as similar CBF in the sham and TBI animals. Medial perilesional (retrosplenial) cortex also had significant reductions in CBF across time (Fig. 3D, Injury, *F*(1,107)=23.0, \*\*\**p*<0.001, Timepoints, *F*(5,107)=3.06, \**p*=0.013). CBF in TBI mice was reduced relative to shams at 3 (\**p*=0.028), 14 (\**p*=0.011), and 60dpi (\**p*=0.021).

**Figure 3.**
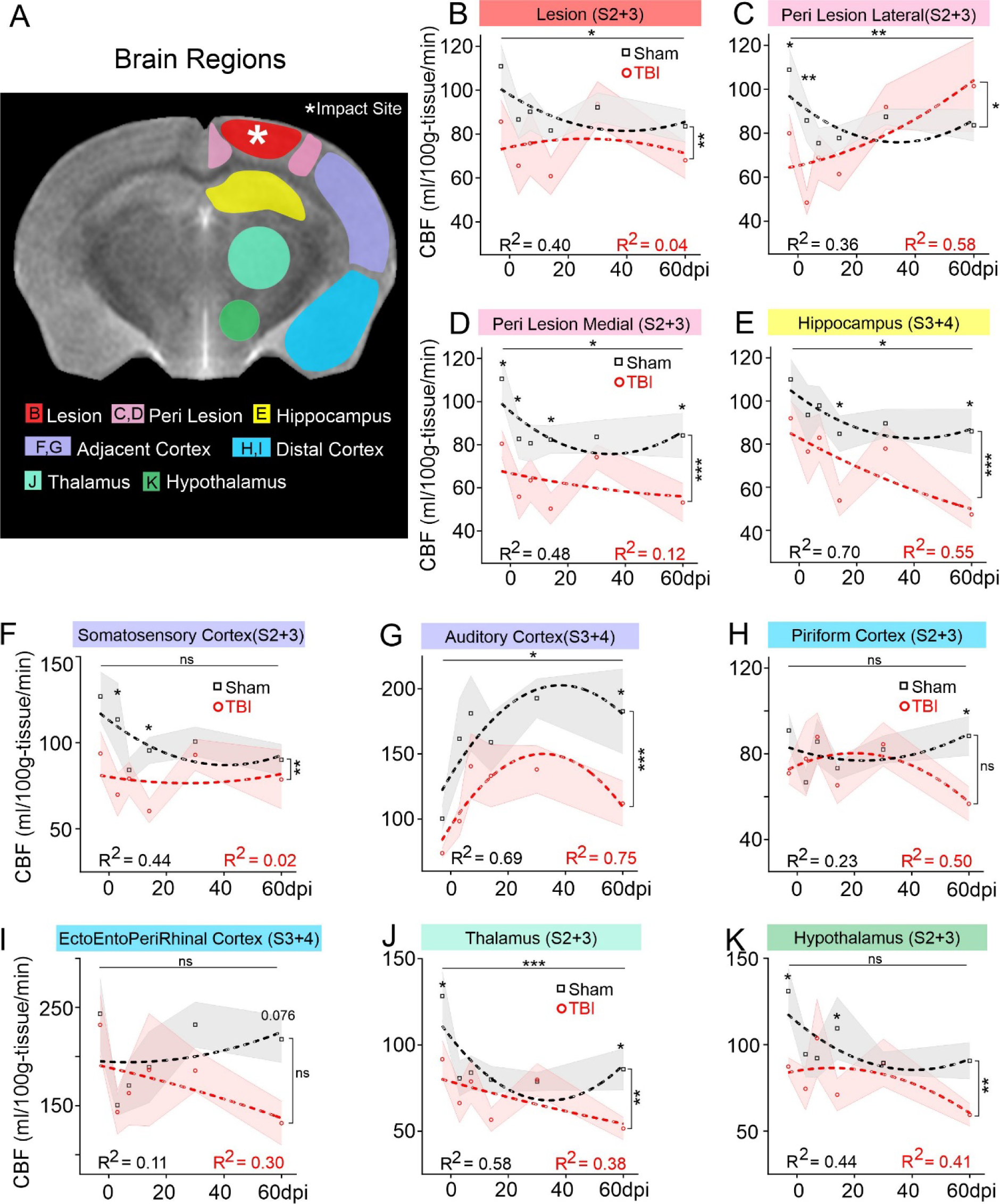
Longitudinal CBF dynamics in brain regions. (A) Representative regions of interest (ROIs) on a coronal MRI**. (B)** CBF at lesion cortex shows overall significant difference across both timepoints (6-sessions 0-60dpi, *F*(5,105)=2.51, \**p*=0.035) and injury condition (Sham vs. TBI, *F*(1,105)=8.36, \*\**p*=0.005). **(C)** Increased CBF in lateral peri-lesional cortex of TBI mice was significant across timepoints (*F*(5,106)=3.77, \*\**p*=0.003) and injury conditions (*F*(1,106)=4.47, \**p*=0.037) and interactions (*F*(5,106)=2.51, \**p*=0.034). **(D)** Medial peri-lesion cortex CBF was significantly reduced across time (*F*(5,107)=3.06, \**p*=0.013) and injury condition (*F*(1,107)=23.0, \*\*\**p*<0.001, interaction–ns). **(E)** CBF profile in dorsal hippocampus was significantly different across time (*F*(5,107)=3.10, \**p*=0.012) and injury condition (*F*(1,107)=14.3, \*\*\**p*<0.001, interaction–ns). **(F)** Somatosensory cortex showed stable CBF across all timepoints (*F*(5,106)=1.88, *p*=0.104, ns) with higher overall longitudinal trend for shams compared to TBI animals (injury factor, *F*(1,106)=10.4, \*\**p*=0.002, interaction–ns). **(G)** Auditory cortex profiles were significantly different across time (*F*(5,107)=3.16, \**p*=0.011) and injury condition (*F*(1,107)=12.7, \*\*\**p*<0.001, interaction–ns). **(H)** Piriform cortex reported stable CBF profiles across timepoints (*F*(5,105)=1.20, *p*=0.316, ns) and injury condition (*F*(1,105)=1.86, *p*=0.175, ns and post-hoc comparisons identified significantly lower CBF at 60dpi in TBI vs sham mice(\**p*=0.025). **(I)** Rhinal cortices (ento, ecto, peri) showed an overall similar CBF trend across time (*F*(5,106)=2.11, *p*=0.069, trending) and injury condition (*F*(1,106)=2.13, *p*=0.147, ns). Post-hoc comparison found a trending decline at 60dpi for TBI vs sham (*p*=0.076). **(J)** CBF in thalamus was significantly different across timepoints (*F*(5,104)=4.57, \*\*\**p*<0.001) and injury condition (*F*(1,104)= 10.0, \*\**p*=0.002, interaction–ns). **(K)** Hypothalamus found stable CBF profiles across time (*F*(5,103)=1.56, *p*=0.177) with significantly different perfusion across injury conditions (*F*(1,103)=7.73, \*\**p*=0.006). (numbers in regional titles denote the PWI slice data were extracted from)

Hippocampus exhibited reduced CBF across the 60dpi period in TBI mice (Fig. 3E, Injury, *F*(1,106)=14.3, \*\*\**p*<0.001, Timepoints, *F*(5,106)=3.10, \**p*=0.012) with significant reductions compared to shams at 14 (\**p*=0.037) and 60dpi (\**p*=0.016). CBF within the cortical regions adjacent to injury (somatosensory, auditory) showed complementary temporal dynamics. In somatosensory cortex TBI mice had reduced CBF spanning the entire experimental period whilst shams had temporal reductions but were not significantly different albeit there was a significant group effect (Fig. 3F, Injury, *F*(1,106)=10.40, \*\**p*=0.002, Timepoints, *F*(5,106)=1.88, *p*=0.104). TBI CBF was significantly lower relative to shams at 3 (\**p*=0.012) and 14dpi (\**p*=0.04). In the adjacent auditory cortex but more distant from TBI site, CBF increased over the first 30dpi but then declined at 60dpi (Fig. 3G, Injury, *F*(1,104)=9.31, \*\**p*=0.003, Timepoints, *F*(5,104)=2.63, \**p*=0.028) with significantly reduced CBF in TBI mice only at 60dpi (\**p*=0.035).

Even more distant from the TBI site, two ventrolateral cortical regions (piriform, ento-ecto- peri-rhinal) exhibited similar declines at 60dpi (Fig. 3H, I), with no significant injury vs time interactions. Piriform cortex was significantly reduced at 60dpi (Post-hoc, Sham vs TBI \**p*=0.025) with trending reductions in the rhinal cortices (ento, ecto, peri) at 60dpi (Post-hoc, Sham vs TBI, *p*=0.076).

Subcortical regions such as thalamus and hypothalamus are involved in social behavior (Fig. 3J, K).^46^ Sham animals showed higher thalamic CBF than TBI mice at baseline (Fig. 3J, Injury, *F*(1,104)=10.0, \*\**p*=0.002, Timepoints, *F*(5,104)=4.57, \*\*\**p*<0.001, *Post hoc*, baseline \**p*=0.013, 60dpi \**p*=0.027). Hypothalamus had no temporally significant CBF changes (*F*(5,103)=1.56, *p*=0.177) but significant differences across injury conditions (*F*(1,103)=7.73, \*\**p*=0.006) with TBI mice having lower CBF at baseline (\**p*=0.017) and 14dpi (\**p*=0.032) compared to shams.

### Reduced cortical angioarchitecture coincides with social behavior decrements at 60dpi

We previously reported recovery of cortical vasculature by 30dpi after TBI.^25^ Surprisingly, at 60pdi we observed broad vascular decrements that mirrored CBF reductions (Fig. 4). At TBI site (lesion), vessel junctions had a trending reduction (*p*=0.068) in TBI mice compared to shams (Fig. 4B), while vessel density was significantly decreased (*p*=0.002, Fig. 4C) accompanied by significantly increased vascular endpoints (*p*=0.0001, Fig. 4D). Average vessel length was unaltered between sham and TBI mice (Fig. 4E). Fractal analysis confirmed reduced vascular complexity that mirrored reduced vessel density (Fig 2F, G). Maximum local fractal dimension (LFD) was significantly reduced at the lesion (*p*=0.0001) and the peri-lesion sites (*p*=0.047) compared to shams (Fig. 4H, 2-tailed-Mann-Whitney Test). Thus, impaired angioarchitecture at 60dpi provides an anatomical basis for our observed physiological and behavioral decrements.

**Figure 4.**
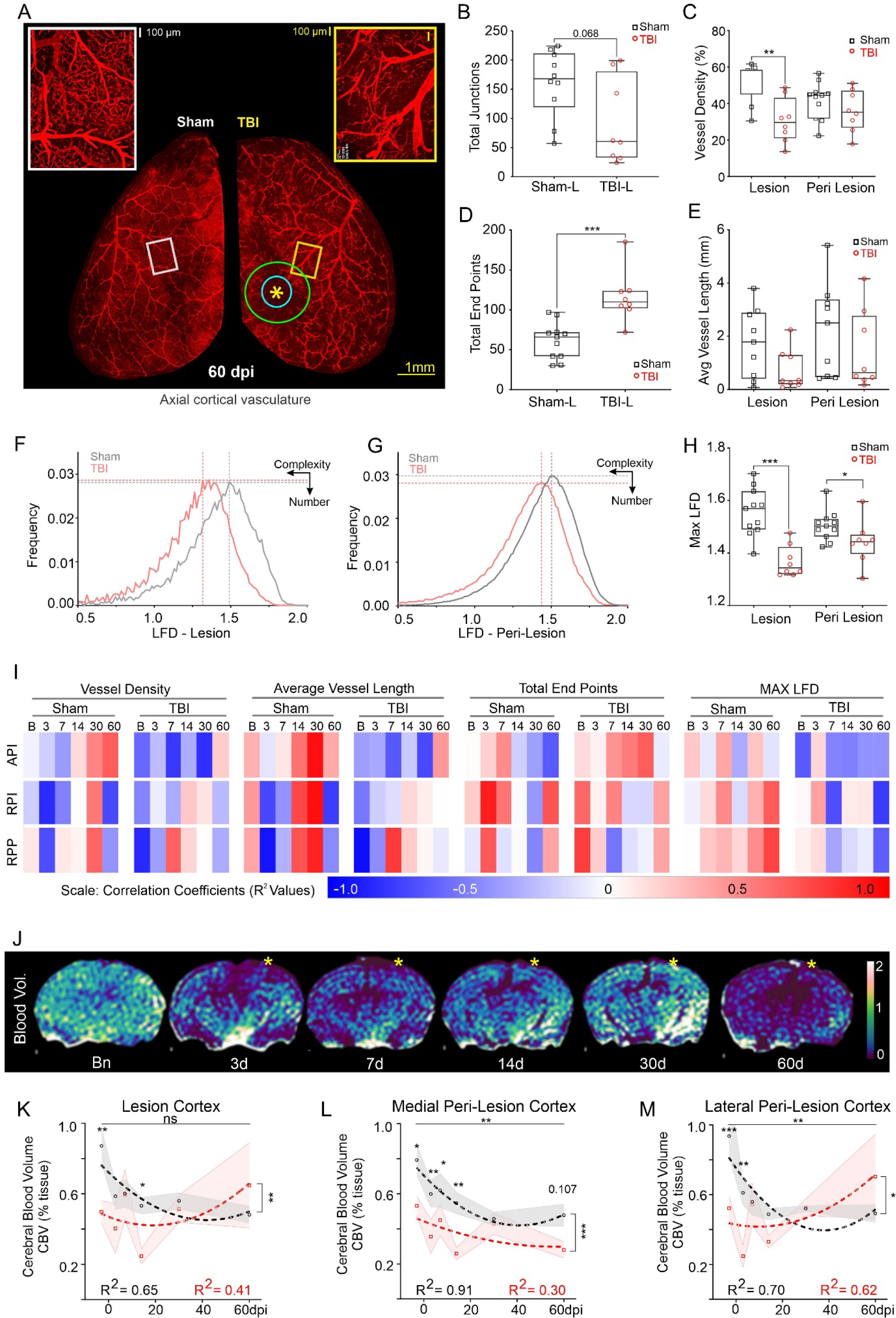
Vascular networks at 60dpi are perturbed. (A) Axial cerebral vasculature is reduced in TBI (right) compared to sham (left) mice in vessel networks encompassing the middle cerebral artery (MCA). Green circle = peri lesion ROI, blue circle = lesion ROI, yellow asterisk = impact site. **(B)** Number of vessel junctions were reduced within the TBI lesion (*p*=0.068) (red circles) compared to shams (black squares). **(C)** Lesion site vessel density in TBI mice was significantly (\*\**p*=0.002) reduced compared to shams but not in peri-lesional cortex. **(D)** Vessel end points were significantly increased at TBI lesion site compared to shams (\*\*\**p*=0.0001). **(E)** Average vessel length was unaltered between sham and TBI mice. **(F, G)** Fractal analysis identified a leftward shift (reduced vessel complexity) in local fractal dimension (LFD) histograms in the lesion **(F)** and in peri-lesion **(G)** sites. **(H)** Maximum local fractal dimension (LFD) was significantly reduced in lesion (\*\*\**p*=0.0001) and peri-lesion sites (\**p*=0.047) for TBI mice relative to shams (B-H: 2-tailed-Mann-Whitney-Test). **(I)** Vascular network parameters at 60dpi were correlated to temporal social outcomes across sham and TBI mice, with opposite correlations between groups. Top row – Absolute partner-interaction-time (API), middle row – Relative partner-interaction time (RPI), bottom row – Relative partner-preference (RPP). **(J)** Transient decrements in cerebral blood volume (CBV, % tissue) over the first 14dpi recovers by 30dpi but is followed by a dramatic 60dpi decrease. **(K)** Lesion cortical CBV was low initially but slowly increased after 30dpi (*F*(5,104)=2.20, *p*=0.059). CBV profiles were significantly different across time but not injury condition (sham vs TBI, *F*(1,104)=7.27, \*\**p*=0.008, interaction–ns). **(L)** Medial peri-lesion cortex had significantly reduced CBV across timepoints (*F*(5,102)=3.76, \*\**p*=0.004) and injury condition (*F*(1,102)=24.2, \*\*\**p*<0.001, interaction–ns). **(M)** CBV in lateral peri-lesion cortex has a similar trajectory as lesion cortex with significant differences across temporal (*F*(5,101)=3.68, \*\**p*=0.004) and injury condition (*F*(1,101)=6.87, \**p*=0.010), and interactions (*F*(5,101)=3.54, \*\**p*=0.005). TBI animals had higher CBV at 60dpi (\*\**p*=0.002).

Further linkage vascular anatomy and longitudinal social behavior was assessed via correlations. Vascular metrics at 60dpi were correlated across temporal social behavior (Fig. 4I). Broadly, sham animals had progressively positive correlations between vessel density and average vessel length against absolute partner-interaction (API) across the 60dpi time course but were negatively correlated in TBI mice (Fig. 4I). Total vessel endpoints were strongly correlated (i.e. more vascular fragmentation) with API while maxLFD negatively correlated (Fig 4I, top row). Both RPI (manual) and RPP (automated) were identical with opposite correlations being observed between groups (Fig. 4I, middle, bottom rows). Interestingly, at 7- and 60dpi, maxLFD had positive correlation with relative partner-interaction for sham but negative correlation with TBI. In sum, TBI mice had opposite correlation trends compared to shams, especially at 60dpi, and the correlation trends across time would suggest that TBI-induced social behavior deficits soon after the injury have predictive potential for long-term vascular decrements.

### Cerebrovascular volumes (CBV) mirror CBF

Cerebral blood volume (CBV) measurements exhibited early post injury declines spanning 14dpi with latent recovery at 30dpi followed by a robust decline at 60dpi (Fig. 4J). CBV in the lesion cortex across time points showed a trending change (*F*(5,104)=2.20, *p*=0.059) and were significantly different across injury conditions (sham vs TBI, *F*(1,104)=7.27, \*\**p*=0.008, interaction – ns) (Fig. 4K). Shams had higher CBV compared to TBI at baseline (\*\**p*=0.006) and at 14dpi (\**p*=0.033). The medial perilesion cortex exhibited significantly different CBV across time (*F*(5,102)=3.76, \*\**p*=0.004) and injury (*F*(1,102)=24.2, \*\*\**p*<0.001, interaction – ns) with elevated CBV in sham mice compared to TBI at baseline (\**p*=0.015), 3 (\*\**p*=0.007), 7 (\**p*=0.048), and 14dpi (\*\**p*=0.005) (Fig. 4L). Baseline CBV for sham animals was higher compared to 30dpi (\**p*=0.044).

CBV in the lateral perilesional cortex (Fig. 4M) was significantly different across timepoints (*F*(5,101)=3.68, \*\**p*=0.004) and injury conditions (*F*(1,101)=6.87, \**p*=0.010) with significant interactions (*F*(5,101)=3.54, \*\**p*=0.005). CBV was initially lower for TBI mice vs. shams at baseline (\*\*\**p*<0.001) and 3dpi (\*\**p*=0.003) but progressively increased whereas, CBV decreased with time in sham mice (Baseline vs. 3dpi: \**p*=0.047, 7dpi: \**p*=0.013, 14dpi: \*\**p*=0.002, 30dpi: \*\**p*=0.008, 60dpi: \*\**p*=0.002). At 60dpi TBI mice had elevated CBV compared to 3dpi (\*\**p*=0.002) indicating persistent CBV increases after injury.

### Spatiotemporally dispersed effects of TBI

The relationship between ipsi- and contralateral brain regions and their CBF was assessed for temporally related patterns (auto-correlation) and interactions between region and CBF (cross-correlations) (Fig. 5A). Broadly, TBI at 3dpi resulted in lower autocorrelations of CBF to ipsilateral compared to contralateral brain regions (Fig. 5A, top panel), which contrasts to the uniform bilateral correlations in shams. Early ipsilateral CBF dysregulation in TBI animals recovered by 30dpi (Fig. 5A, middle panel), which coincides with vascular recovery.^25^ However, at 60dpi when both CBF and vessel density are reduced, CBF auto-correlations across multiple brain regions are dramatically reduced (Fig. 5A, bottom panel) in stark contrast to sham mice that exhibit strong bilateral CBF auto-correlations, as would be expected in healthy mice. These findings confirm the prolonged secondary consequences of TBI on blood flow across broad portions of the brain, including those distant from the injury and mirror angioarchitecture.

**Figure 5.**
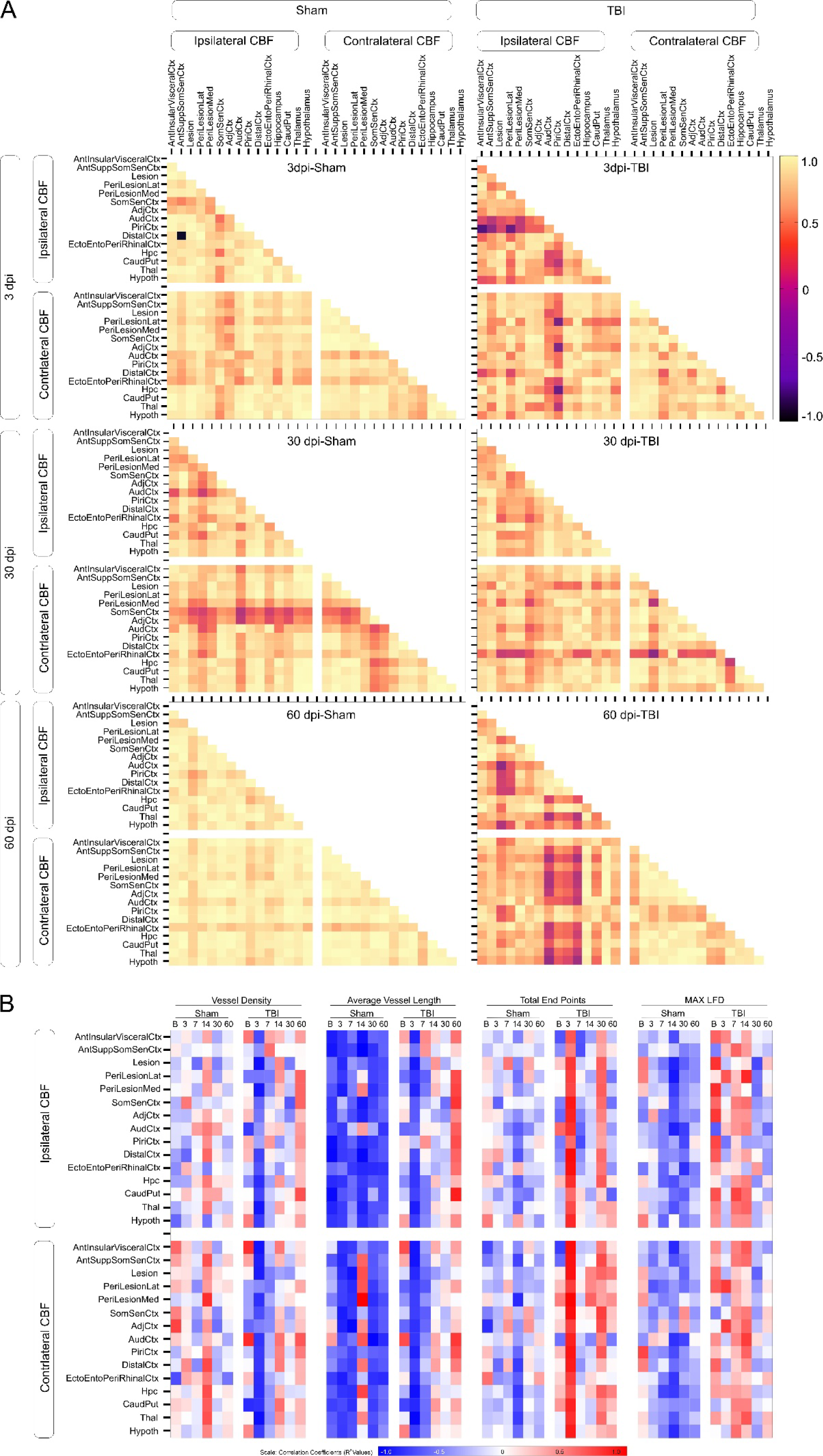
Temporal relationships between cerebral blood flow (CBF) across brain regions. (A) Temporal CBF correlation coefficients across brain regions highlight global alterations due to TBI (right panel) resulting in loss of CBF auto-correlations at the injury site at 3dpi that moderately recovers by 30dpi but is greatly perturbed at 60dpi unlike shams (left panel). This dysregulation also spreads to the contralateral hemisphere at 60dpi in TBI mice. **(B)** Temporal CBF correlations to vascular network measures at 60dpi further confirm an initial recovery. However, delayed cerebrovascular structural deficits contributes to the declining brain perfusion. At 60dpi, vessel density in lesion cortex and longitudinal CBF across ipsi- and contralateral brain regions in TBI animals show mostly negative correlations at 3-30dpi followed by positive correlations at 60dpi. In contrast, sham animals show highly positive correlations at 14dpi and low mixed correlations at other time points. Vessel length vs. CBF correlations are negative for sham animals across all timepoints but positive for TBI animals at 60dpi. Total end points and CBF correlations in sham animals also exhibited mostly mixed correlations except 14dpi with negative correlations. Conversely, TBI animals show positive correlations at 3- and 30dpi, negative ipsilateral correlations for 7-60dpi, and negative contralateral correlations at 7dpi, but positive contralateral CBF correlations at 14-60dpi. maxLFD and CBF correlations were negative for sham animals unlike TBI animals with positive correlations at 7-14dpi, negative at 30dpi, and positive again at 60dpi. Abbreviations: Med – Medial, Lat – Lateral, Ctx – Cortex, Hpc – Hippocampus, CaudPut – Caudate putamen, Thal – Thalamus, Hypoth - Hypothalamus

We next probed if early CBF dynamics (3-30dpi) predict the imminent secondary vascular damage late after injury (60dpi). Correlations between longitudinal CBF (Baseline–60dpi) and vascular metrics measured within lesion cortex at 60dpi (Fig. 5B) demonstrate distinct longitudinal correlation patterns in sham and TBI mice. Roughly, in TBI mice the correlations suggest an initial negative correlation(s) between vessel density and length that increasingly, with time become strongly correlated by 60dpi. These observations are opposite in vessel endpoints and vascular complexity measures (LFD). Thus, early (3-7dpi) blood flow and vascular disruption are not synchronized whereas the low CBF and loss of the vascular network are tightly correlated at 60dpi. In summary, CBF measures after TBI may reflect altered vascular morphology.

### Neurovascular function corresponds to social behavior decrements

To capture associations between cerebrovascular function (CBF, CBV) and social behavior longitudinally after TBI, we undertook correlations across bilateral regions. This approach demonstrated strong linkage of RPI to ipsi- and contralateral cerebrovascular decrements across social behavior related brain regions (ipsilateral, Fig. 6A-C; contralateral, Fig. 6G-I). In general, the correlation matrices, as evident in similar heatmaps between ispi- and contralateral regions were remarkably consistent. When all regional correlations were averaged we observed distinct signatures that separated TBI mice from shams (Fig. 6B-F, H-L). TBI induced ipsilateral CBF and CBV changes strongly which correlated with RPI and API but was negatively correlated in sham mice. Compared to CBF, the CBV changes exhibited the strongest correlations on both ipsi- and contralateral brain regions (e.g. Fig. 6C, I). In summary, CBF vs. RPI/API correlations increase from 3- 30dpi but then decline by 60dpi while sham animals show opposite trajectories (Fig. 6B, E). CBV correlation profiles with RPI/API have progressive negative correlations in shams whereas TBI mice have persistent positive correlations (Fig. 6C, F). Similar correlational directionality was observed between CBF/CBV and RPI/API in contralateral brain regions (Fig. 6 G-L). Profiles showing API correlated with ipsilateral (Fig. 6D-F) and contralateral (Fig. 6J-L) Hence, early correlations between behavior and CBF predict long-term physiological deficits.

**Figure 6.**
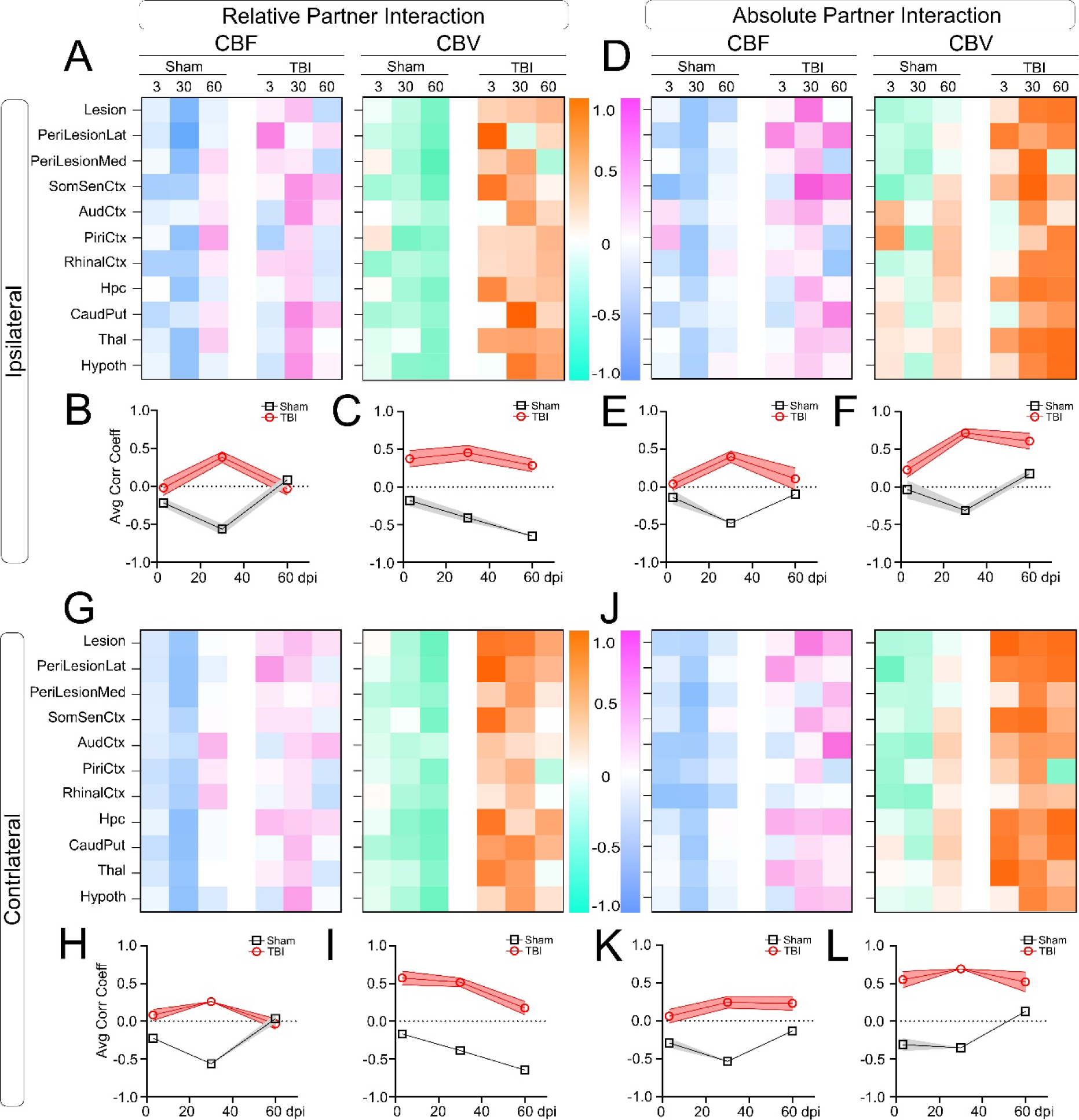
Neurovascular physiology in social behavior associated brain regions . (A-C) Correlation coefficients of relative partner interaction (RPI) to CBF and CBV across ipsilateral social behavior associated brain regions in TBI and sham mice at 3, 30, and 60dpi (columns). Regionally averaged correlation coefficients are illustrated in B, C. **(B)** TBI animals exhibit the opposite correlations at 30dpi with strong positive correlations unlike negative correlations in shams for RPI vs. CBF. **(C)** In TBI mice RPI vs. CBV show strong correlations across time in contrast to shams having negative correlations with behavior. **(D)** Correlations of absolute partner interaction (API) with ipsilateral CBF and CBV exhibit a virtually identical (compared to RPI) set of regional correlations between sham and TBI animals. **(E)** API vs. CBF correlations exhibit identical trends as RPI vs. CBF correlations for both sham and TBI animals with the largest divergence at 30dpi. **(F)** API vs. CBV correlations are elevated in TBI mice compared to reduced strength correlation patterns in shams. **(G)** Correlation coefficients of RPI vs. CBF and CBV across contralateral brain regions behavior have the same temporal patterns as the ipsilateral brain regions. **(H)** Contralateral CBF vs. RPI at 30dpi have increased correlations in TBI mice but reduced in shams with no overt differences at 3 or 60dpi. **(I)** As in the ipsilateral regions, RPI vs. CBV correlations were strongly positive in TBI mice but greatly reduced temporally in shams **(J)** API correlation to contralateral CBF and CBV were strong across all regions in TBI but not shams. **(K)** API vs. CBF correlations in shams were negative compared to TBI animals showing stable moderate positive correlations. **(L)** API vs. CBV correlation patterns were strongly positive for TBI mice but predominately negative in sham animals. Abbreviations: Med – Medial, Lat – Lateral, Ctx – Cortex, Hpc – Hippocampus, CaudPut – Caudate putamen, Thal – Thalamus, Hypoth - Hypothalamus

## Discussion

We investigated longitudinal and dynamic evolution of behavior and vascular physiology across 2-months post-TBI (Fig. 1). Our study is the first preclinical TBI study to demonstrate reduced social preference between familiar cage-mates when vascular functions decline. The key findings are: i) a late decline (60dpi) in sociability of TBI animals while motor and exploratory behavior recover; ii) longitudinal CBF (Figs. 2, 3) identifying a novel multiphasic recovery profile with delayed 60dpi global decline in perfusion that also encompasses brain regions processing social behavior; iii) angioarchitecture exhibits vascular damage at 60dpi, including reduced vessel density and increased fragmentation (Fig. 4); iv) vascular and social behavior correlations displayed pronounced negative correlations early after TBI but progressively become positively correlated by 60dpi; iv) regional correlations confirmed dynamic bilateral CBF changes across the brain after a unilateral TBI, (Fig. 5) that becomes predominately perturbed ipsilaterally; and v) unique correlation patterns between behavior and vascular physiology emerge in TBI animals (Fig. 6). In summary, our study for the first time links how cerebrovascular physiology contributes to decline in social interactions after TBI in male mice. Our findings also suggest that monitoring social behavior early after brain injury may predict long-term neurovascular damage, providing a putative avenue for therapeutic interventions.

Neuropsychological disabilities, including sociability issues are being increasingly recognized after TBI in adults^47^ and children.^48^ Subjects often exhibit the inability to recognize affective facial expressions,^49, 50^ impaired verbal and non-verbal communication,^9^ and diminished empathy,^51^ culminating into decreased time spent with friends and families. Interestingly, brain regions regulating social behavior are common between rodents and primates.^52, 53^ Feature content of socially transferred food preference memory is known to decline after systems consolidation,^40^ and social isolation elicits similar pathophysiology in healthy rodents as in humans.^54^ A few studies, in preclinical models of TBI, have explored social dominance and interaction behavior.^20, 55^ Our observation of behavioral recovery at 30dpi corroborates a recent study that showed lack of social deficits at 30dpi in mice with repeated mTBI.^56^ Consistent with the previous literature we also found TBI animals cover longer distances than the shams, at higher relative speed.^57^ Preferentially exploring and frequenting the center of the open-field arena suggests an inclination for taking risks, a well-documented behavior observed in rodents and human subjects after TBI.^58–60^ However, long-term social isolation among familiar individuals post-TBI has not been studied.

In chronic TBI, cognitive impairments and social isolation often precede the debilitating prolonged consequences of injury among human subjects that parallel Alzheimer’s-related signatures,^61^ depression, and suicidal tendencies.^62^ Our novel approach captures the long-term social interaction deficits between familiar mice as observed in head-injured human subjects.^63, 64^ In the classic Crawley’s 3-chamber test,^37, 65^ a test mouse interacts with a combination of unfamiliar mice and a novel object where the social performance reflects combined effect of novel entity and neophobia.^66^ An adult frontal TBI model showed increased preference for familiar mice in 3-chamber task, however, longitudinal transition of social behavior in same animals was not investigated.^16^ Our temporal study demonstrated, in TBI mice, no initial sociability deficits but at chronic time points the emergence of a decline in voluntary preference of a known animal i.e. their cage-mate partner. This decrement in sociability of TBI mice was accompanied by decreased anxiety and increased exploration consistent with increased risk behavior, as previously reported.^67^

Numerous studies have sought to link altered behavior to inflammation^68^ and neuronal cell death^69^ as underlying mechanisms. Surprisingly, there are limited investigations into how cerebrovascular morphology and function are linked after TBI to behavioral deficits. A recent study in TBI subjects found significant associations at ∼2yrs post injury between decreased perfusion and psychoemotional outcomes (i.e. anxiety).^70^ Neurovascular coupling implies a link between task-evoked cellular metabolic demands and blood flow to the activated brain region.^71^ Social recognition associated cellular networks require protein synthesis and includes cAMP responsive element-binding protein (CREB) for transcriptional consolidation.^72, 73^ Adult human TBI leads to resting hypoperfusion in many task-related brain regions years after injury and is similar to that reported in TBI rodent models.^31, 74^ In our study we also found early hypoperfusion that recovered by 30dpi but then rapidly declined in virtually every brain region. Interestingly at 60 dpi, we find reduced CBF in dorsal hippocampal (dHpC) whose subregions are necessary for spatial exploration (CA1) and social interaction (CA2).^75^. Reduction in interaction with familiar in comparison to novel conspecifics in chronic-TBI could be driven by corticotrophin signaling from PFC to lateral septum.^76^ Two months after the injury, reduction in intrinsic excitability and decreased synaptic output of somatostatin neurons in layer V of orbito-frontal cortex has been reported^77^, consistent with long-term behavioral deficits as shown in ours and other studies.^78, 79^ could arise from the TBI-associated neuronal damage in cortex, dorsal hippocampus and thalamus Future studies can shed more light on the neural underpinnings of similar TBI comorbidities. Along this rationale, our data captures a delayed decline in overall neurovascular functional health and maintenance in brain regions associated with social behavior. Our results show reduced regional CBF post TBI with a novel multiphasic pattern of intermittent recovery at 7 and 30dpi, worsening of blood flow at 14dpi and a delayed decline at 60dpi. Interestingly, we observe similar longitudinal profiles for RPP and CBF measurements in social behavior relevant brain regions.

Our results are consistent with previous studies showing acute and chronic hypoperfusion and behavior deficits in both clinical and preclinical TBI.^57, 80–82^ Several studies have demonstrated recovery of behavior performance after TBI in rats, ^83^ mice, ^57, 84^ and humans.^85^ The primary finding of many studies is that social, anxiety and cognitive behavioral domains worsen with increasing time. A similar progressive decline in brain perfusion long-term has been reported. Human studies report CBF recovery by 3 weeks post injury, with linkage to improved neurological outcomes.^86^ Grohn and colleagues noted biphasic hypoperfusion with transient recovery followed by a second hypoperfusion epoch over two weeks after TBI, that relates to changes in vascular density.^87^ Others have also noted protracted CBF reduction in rodents has been shown to last up to 1 year post TBI.^88^ Our current study and previous observations demonstrated perfusion recovery at 30dpi ^25^ that was reflected by recovery of vascular density. In our study the consistent decrement in CBF at 60dpi relates to reduced vascular density and complexity which thereby exacerbates behavioral deficits.

In support of our study, previous findings noted focal TBI elicits longitudinal global cerebrovascular deficits underlying large-scale brain network effects, likely leading to protracted social deficits.^89^ Bilateral CBF reductions, similar to our observations, have been seen in human mTBI subjects for prefrontal cortex, putamen, and hippocampus, while reduced CBF in cortex and caudate putamen is associated with depressive symptoms, and in hippocampus with anxiety.^70^ Consistent with the previous human studies, we also observe thalamic pathophysiology after mTBI.^90^ Decreased thalamic dendrite complexity in rats also shows recovery by 4-weeks post mTBI,^91^ corroborated by corresponding vascular and functional perfusion recovery in our study that then to collapses by 60dpi.

TBI results in immediate damage to focal and distant cerebrovascular morphology that then partially recovers.^23, 25, 92^ The subsequent secondary cellular and molecular cascades after moderate to severe TBI result in long-term deficits including hemorrhage, edema, reduced CBF, vasospasms, blood-brain disruptions, coagulopathy, and chronic inflammation.^93–95^

Surprisingly, we observed a second period of vascular loss at 60dpi despite vascular recovery by 30dpi.^25^ The late diminished vessel characteristics mirrored reduced CBF virtually across all brain regions compared to shams. Thus, initial vascular recovery is transient and is not well integrated within the brain parenchyma as stable neurovascular units.^96, 97^ Structural vascular abnormalities in human TBI subjects exhibit microvessels with flattened, reduced lumina and longitudinal folds in the pial, cortical, and capillary zones.^26, 98^ In lateral fluid percussion rodent models there also is microvascular recovery which does not mirror healthy control vasculature.^99^ ^87^ As noted previously, we observed identical recovery profiles after TBI^25^ which then rapidly degrade by 60dpi. Vascular density was also increased at 14dpi after repeated TBI concomitant with diminished CBF, cerebrovascular reactivity, and neuronal activity.^100^ In adult and pediatric human subjects after TBI, CBF and CBV decline.^47, 101^ Broadly, TBI in clinical and preclinical studies suggest that dynamic vascular density alterations lead to chronic reductions in brain responsivity and perfusion.

It is noteworthy that there are regional variations, particularly in corelative preclinical studies. For example, hippocampal vessel density does not vary with declining CBF whereas increases in CBF and vessel density were reported in ipsilateral thalamus 8-months after TBI.^28, 31, 47^ The authors reported that poor spatial exploration performance correlated with increased thalamic vessel density. Griffiths and colleagues reported no changes in cortical or hippocampal CBF or CBV 6mo after mild TBI despite cognitive decrements.^102^ Similar findings have been reported in individuals with mild^70^and in moderate severe TBI.^103^

Our correlations measure the interdependent blood flow across brain regions highlighting bilateral effects specific to TBI animals. Specifically, global blood flow correlations were reduced at 3dpi with recovery by 30dpi followed by a dramatic decline at 60dpi. The correlations of longitudinal behavior to 60dpi vascular metrics provided an early-stage behavioral marker to predict the imminent long term neurovascular damage from TBI. Finally, our correlations between social behavior, blood flow and blood volume in brain regions exhibited unique patterns for TBI animals at 3, 30- and 60dpi. Similar behavioral correlations to CBF were found in mTBI.^102^

The strengths of our study are, i) longitudinal in vivo assessments of behavior alongside cerebrovascular function after TBI which provide a continuous view of how recovery is modulated; ii) behavioral tests across multiple domains (motor, exploratory, social) but with novel social preference for familiar cage mates, an observation reported in human subjects yet underexplored in preclinical models; iii) our extended observation window to 2months post-TBI which is equivalent to ∼7 years in humans^104^, iv) novel multiphasic global evolution of brain perfusion after injury in the same subjects; v) novel CBF autocorrelations across whole brain regions demonstrating remission of cerebrovascular pathophysiology at 30dpi but recurrence at 60dpi; vi) the loss of structural vascular networks underlie CBF and social behavior deficits; and vi) correlations across behavior and cerebrovascular physiology providing a predictive assessment of imminent long-term pathophysiology underlying TBI comorbidities.

There are several limitations of our current study. They include: i) absence of female mice perfusion and behavioral data. Female gender is underrepresented in clinical and most pre- clinical research studies and emerging studies suggest that women of the same age group (compared to men) are more susceptible to adverse consequences of TBI.^105^ To fill this gap we are currently investigating the long term physiological and behavioral pathology post-TBI in female mice; ii) some of the variance in our measurements can be attributed to the modest number of replicates (n=6-8/grp/time) but exhibited sufficient statistical power particularly in light of our longitudinal assessments; iii) limited anatomical resolution from the perfusion-weighted MRI measurements. Due to the rapid acquisition techniques in-plane resolution was 250um/pixel which provide sufficient resolution for regional brain assessments using manual segmentations based on the Allen brain atlas^106^. Future studies will address these limitations by combining high-resolution optical with magnetic resonance imaging. MRI does provide a powerful non-invasive and longitudinal global assessment of pathophysiology that is not feasible with other techniques; and iv) baseline CBF variations between sham and TBI groups before injury were noted in some but were not different in regions involved in social behavior (hippocampus, piriform, auditory, rhinal cortices).

## Conclusions and Future Directions

In conclusion, our study for the first time demonstrates the cerebrovascular underpinnings of emerging social behavioral deficits after chronic TBI. Social interactions among familiar mice long after a TBI were reduced with concurrent longitudinal physiological reductions in CBF and CBV. A steep decline in CBF at 60dpi in social behavior related brain regions, was observed in hippocampus and rhinal cortex. The loss of angioarchitecture at 60dpi provides the basis for precipitous declines in CBF and social behavior. Further, our correlations point to broad linkage between impairments in vascular metrics, CBF, CBV, and social behavior metrics. We suggest that such correlations may have predictive value for obtaining early estimates of long-term damage, and potentially informing the optimal treatments. In addition to pharmacological interventions, enriched environments with monitored exercise^107–109^ and virtual environments are deemed helpful during chronic TBI recovery in rodents^110^ and humans ^111, 112^, including virtual social networks.^113^ Future investigations, in addition to assessing influence of sex should investigate how vascular smooth muscle attributes are modified by TBI ^114^ and how pericytes regulate blood flow.^115^ Finally, chronic TBI sequelae such as BBB dysfunction, TGFβ signaling, and neuroinflammation also contribute to the long-term sequelae injury. While our study in a rodent preclinical model hints at the linkage between neuropsychological outcomes modulated by brain perfusion, continued investigations are needed to improve our understanding of the longitudinal implications TBI and how we might best intervene to improve patient outcomes.

## Data availability

Data is available at a reasonable request from the corresponding author.

## Acknowledgements

We thank Dr. Amandine Jullienne for training on tail vein injections and vessel painting, Rojina Pad for help with imaging vessel painted brains, Brenda Noarbe for help with implementing MRI analysis routines, and UCI ULAR staff for taking care of the animals.

## Supplementary material

Supplementary material is available.

## Author Contributions

AS – Conceptualization, Behavior and MRI - experiment design and execution, vessel painting, microcopy imaging, data processing, analyses, and visualization, manuscript writing and editing

SG – Behavior experiment execution, MRI data processing, microcopy imaging

AV – Optical imaging, Data processing for semi-automated behavior and microcopy imaging SL – MRI data collection

AO – Funding acquisition, conceptualization, MRI sequence design, MRI experiment design and execution, data processing and analyses, manuscript writing and editing

## References

1. Dewan MC, Rattani A, Gupta S, et al. Estimating the global incidence of traumatic brain injury. J Neurosurg. Apr 27 2018;130(4):1080–1097. doi:10.3171/2017.10.JNS17352

2. Blaya MO, Raval AP, Bramlett HM. Traumatic brain injury in women across lifespan. Neurobiol Dis. Mar 2022;164:105613. doi:10.1016/j.nbd.2022.105613

3. Bailey DM, Jones DW, Sinnott A, et al. Impaired cerebral haemodynamic function associated with chronic traumatic brain injury in professional boxers. Clin Sci (Lond). Feb 2013;124(3):177–89. doi:10.1042/CS20120259

4. McKee AC, Robinson ME. Military-related traumatic brain injury and neurodegeneration. Alzheimers Dement. Jun 2014;10(3 Suppl):S242-53. doi:10.1016/j.jalz.2014.04.003

5. Li LM, Carson A, Dams-O’Connor K. Psychiatric sequelae of traumatic brain injury - future directions in research. Nat Rev Neurol. Sep 2023;19(9):556–571. doi:10.1038/s41582-023-00853-8

6. McKee AC, Daneshvar DH. The neuropathology of traumatic brain injury. Handb Clin Neurol. 2015;127:45–66. doi:10.1016/B978-0-444-52892-6.00004-0

7. Obenaus A. Traumatic brain injury. Encyclopedia of Mental Health. 2023:519–534.

8. Chen F, Chi J, Niu F, et al. Prevalence of suicidal ideation and suicide attempt among patients with traumatic brain injury: A meta-analysis. J Affect Disord. Mar 1 2022;300:349–357. doi:10.1016/j.jad.2022.01.024

9. Rousseaux M, Verigneaux C, Kozlowski O. An analysis of communication in conversation after severe traumatic brain injury. Eur J Neurol. Jul 2010;17(7):922–9. doi:10.1111/j.1468-1331.2009.02945.x

10. Hoofien D, Gilboa A, Vakil E, Donovick PJ. Traumatic brain injury (TBI) 10-20 years later: a comprehensive outcome study of psychiatric symptomatology, cognitive abilities and psychosocial functioning. Brain Inj. Mar 2001;15(3):189–209. doi:10.1080/026990501300005659

11. Rigon A, Duff MC, Beadle J. Lonely But Not Alone: Neuroticism Mediates the Relationship Between Social Network Size and Loneliness in Individuals With Traumatic Brain Injury. J Int Neuropsychol Soc. Mar 2019;25(3):285–292. doi:10.1017/S1355617718001108

12. Andrews TK, Rose FD, Johnson DA. Social and behavioural effects of traumatic brain injury in children. Brain Inj. Feb 1998;12(2):133–8. doi:10.1080/026990598122755

13. Khan N, Ryan NP, Crossley L, Hearps S, Catroppa C, Anderson V. Associations Between Peer Relationships and Self-Esteem After Childhood Traumatic Brain Injury: Exploring the Mediating Role of Loneliness. J Neurotrauma. Oct 2023;40(19-20):2100–2109. doi:10.1089/neu.2022.0420

14. Sander AM, Struchen MA. Interpersonal relationships and traumatic brain injury. J Head Trauma Rehabil. Jan-Feb 2011;26(1):1–3. doi:10.1097/HTR.0b013e3182068588

15. Salas CE, Casassus M, Rowlands L, Pimm S, Flanagan DAJ. “Relating through sameness”: a qualitative study of friendship and social isolation in chronic traumatic brain injury. Neuropsychol Rehabil. Oct 2018;28(7):1161–1178. doi:10.1080/09602011.2016.1247730

16. Chou A, Morganti JM, Rosi S. Frontal Lobe Contusion in Mice Chronically Impairs Prefrontal-Dependent Behavior. PLoS One. 2016;11(3):e0151418. doi:10.1371/journal.pone.0151418

17. Ritzel RM, Li Y, He J, et al. Sustained neuronal and microglial alterations are associated with diverse neurobehavioral dysfunction long after experimental brain injury. Neurobiol Dis. Mar 2020;136:104713. doi:10.1016/j.nbd.2019.104713

18. Nolan A, Hennessy E, Krukowski K, et al. Repeated Mild Head Injury Leads to Wide-Ranging Deficits in Higher-Order Cognitive Functions Associated with the Prefrontal Cortex. J Neurotrauma. Oct 15 2018;35(20):2425–2434. doi:10.1089/neu.2018.5731

19. Algamal M, Saltiel N, Pearson AJ, et al. Impact of Repetitive Mild Traumatic Brain Injury on Behavioral and Hippocampal Deficits in a Mouse Model of Chronic Stress. J Neurotrauma. Sep 1 2019;36(17):2590–2607. doi:10.1089/neu.2018.6314

20. Semple BD, Canchola SA, Noble-Haeusslein LJ. Deficits in social behavior emerge during development after pediatric traumatic brain injury in mice. J Neurotrauma. Nov 20 2012;29(17):2672–83. doi:10.1089/neu.2012.2595

21. Khodaie B, Lotfinia AA, Ahmadi M, et al. Structural and functional effects of social isolation on the hippocampus of rats with traumatic brain injury. Behav Brain Res. Feb 1 2015;278:55–65. doi:10.1016/j.bbr.2014.09.034

22. Doulames VM, Vilcans M, Lee S, Shea TB. Social interaction attenuates the extent of secondary neuronal damage following closed head injury in mice. Front Behav Neurosci. 2015;9:275. doi:10.3389/fnbeh.2015.00275

23. Obenaus A, Ng M, Orantes AM, et al. Traumatic brain injury results in acute rarefication of the vascular network. Sci Rep. Mar 22 2017;7(1):239. doi:10.1038/s41598-017-00161-4

24. Griffin AD, Turtzo LC, Parikh GY, et al. Traumatic microbleeds suggest vascular injury and predict disability in traumatic brain injury. Brain. Nov 1 2019;142(11):3550–3564. doi:10.1093/brain/awz290

25. Lin X, Chen L, Jullienne A, et al. Longitudinal dynamics of microvascular recovery after acquired cortical injury. Acta Neuropathol Commun. Apr 25 2022;10(1):59. doi:10.1186/s40478-022-01361-4

26. Rodriguez-Baeza A, Reina-de la Torre F, Poca A, Marti M, Garnacho A. Morphological features in human cortical brain microvessels after head injury: a three-dimensional and immunocytochemical study. Anat Rec A Discov Mol Cell Evol Biol. Jul 2003;273(1):583–93. doi:10.1002/ar.a.10069

27. Yu GX, Mueller M, Hawkins BE, et al. Traumatic brain injury in vivo and in vitro contributes to cerebral vascular dysfunction through impaired gap junction communication between vascular smooth muscle cells. J Neurotrauma. Apr 15 2014;31(8):739–48. doi:10.1089/neu.2013.3187

28. Amyot F, Kenney K, Spessert E, et al. Assessment of cerebrovascular dysfunction after traumatic brain injury with fMRI and fNIRS. Neuroimage Clin. 2020;25:102086. doi:10.1016/j.nicl.2019.102086

29. Wei EP, Hamm RJ, Baranova AI, Povlishock JT. The long-term microvascular and behavioral consequences of experimental traumatic brain injury after hypothermic intervention. J Neurotrauma. Apr 2009;26(4):527–37. doi:10.1089/neu.2008.0797

30. Xu X, Cowan M, Beraldo F, et al. Repetitive mild traumatic brain injury in mice triggers a slowly developing cascade of long-term and persistent behavioral deficits and pathological changes. Acta Neuropathol Commun. Apr 6 2021;9(1):60. doi:10.1186/s40478-021-01161-2

31. Hayward NM, Immonen R, Tuunanen PI, Ndode-Ekane XE, Grohn O, Pitkanen A. Association of chronic vascular changes with functional outcome after traumatic brain injury in rats. J Neurotrauma. Dec 2010;27(12):2203–19. doi:10.1089/neu.2010.1448

32. Wang Y, Yu S, Li M. Neurovascular crosstalk and cerebrovascular alterations: an underestimated therapeutic target in autism spectrum disorders. Front Cell Neurosci. 2023;17:1226580. doi:10.3389/fncel.2023.1226580

33. Paloyelis Y, Doyle OM, Zelaya FO, et al. A Spatiotemporal Profile of In Vivo Cerebral Blood Flow Changes Following Intranasal Oxytocin in Humans. Biol Psychiatry. Apr 15 2016;79(8):693–705. doi:10.1016/j.biopsych.2014.10.005

34. Knudsen LV, Sheldrick AJ, Vafaee MS, Michel TM. Diversifying autism neuroimaging research: An arterial spin labeling review. Autism. Jul 2023;27(5):1190–1203. doi:10.1177/13623613221137230

35. Salehi A, Jullienne A, Wendel KM, et al. A Novel Technique for Visualizing and Analyzing the Cerebral Vasculature in Rodents. Transl Stroke Res. May 15 2018;doi:10.1007/s12975-018-0632-0

36. Singh A, Balaji J. Sensitive Estimation of Flavor Preferences in STFP Using Cumulative Time Profiles. Bio Protoc. Nov 5 2017;7(21):e2601. doi:10.21769/BioProtoc.2601

37. Crawley JN. Twenty years of discoveries emerging from mouse models of autism. Neurosci Biobehav Rev. Mar 2023;146:105053. doi:10.1016/j.neubiorev.2023.105053

38. Schindelin J, Arganda-Carreras I, Frise E, et al. Fiji: an open-source platform for biological-image analysis. Nat Methods. Jun 28 2012;9(7):676–82. doi:10.1038/nmeth.2019

39. Chiara V, Kim SY. AnimalTA: A highly flexible and easy-to-use program for tracking and analysing animal movement in different environments. Methods in Ecology and Evolution. 2023;14(7):1699–1707. doi:10.1111/2041-210x.14115

40. Singh A, Kumar S, Singh VP, Das A, Balaji J. Flavor Dependent Retention of Remote Food Preference Memory. Front Behav Neurosci. 2017;11:7. doi:10.3389/fnbeh.2017.00007

41. Friard O, Gamba M, Fitzjohn R. BORIS: a free, versatile open-source event-logging software for video/audio coding and live observations. Methods in Ecology and Evolution. 2016;7(11):1325–1330. doi:10.1111/2041-210x.12584

42. Salehi A, Salari S, Jullienne A, et al. Vascular topology and blood flow are acutely impacted by experimental febrile status epilepticus. J Cereb Blood Flow Metab. Jan 2023;43(1):84–98. doi:10.1177/0271678X221117625

43. Parker GJ, Roberts C, Macdonald A, et al. Experimentally-derived functional form for a population-averaged high-temporal-resolution arterial input function for dynamic contrast-enhanced MRI. Magn Reson Med. Nov 2006;56(5):993–1000. doi:10.1002/mrm.21066

44. Petrella JR, Provenzale JM. MR perfusion imaging of the Brain: Techniques and applications. Review. Am J Roentgenol. Jul 2000;175(1):207–219. doi:DOI 10.2214/ajr.175.1.1750207

45. Zudaire E, Gambardella L, Kurcz C, Vermeren S. A computational tool for quantitative analysis of vascular networks. PLoS One. 2011;6(11):e27385. doi:10.1371/journal.pone.0027385

46. Usui N. Possible roles of deep cortical neurons and oligodendrocytes in the neural basis of human sociality. Anat Sci Int. Jan 2024;99(1):34–47. doi:10.1007/s12565-023-00747-1

47. Morton MV, Wehman P. Psychosocial and emotional sequelae of individuals with traumatic brain injury: a literature review and recommendations. Brain Inj. Jan 1995;9(1):81–92. doi:10.3109/02699059509004574

48. Lopez DA, Christensen ZP, Foxe JJ, Ziemer LR, Nicklas PR, Freedman EG. Association between mild traumatic brain injury, brain structure, and mental health outcomes in the Adolescent Brain Cognitive Development Study. Neuroimage. Nov 2022;263:119626. doi:10.1016/j.neuroimage.2022.119626

49. Knox L, Douglas J. Long-term ability to interpret facial expression after traumatic brain injury and its relation to social integration. Brain Cogn. Mar 2009;69(2):442–9. doi:10.1016/j.bandc.2008.09.009

50. Rigon A, Voss MW, Turkstra LS, Mutlu B, Duff MC. Different aspects of facial affect recognition impairment following traumatic brain injury: The role of perceptual and interpretative abilities. J Clin Exp Neuropsychol. Oct 2018;40(8):805–819. doi:10.1080/13803395.2018.1437120

51. de Sousa A, McDonald S, Rushby J, Li S, Dimoska A, James C. Understanding deficits in empathy after traumatic brain injury: The role of affective responsivity. Cortex. May 2011;47(5):526–35. doi:10.1016/j.cortex.2010.02.004

52. Kim Y, Venkataraju KU, Pradhan K, et al. Mapping social behavior-induced brain activation at cellular resolution in the mouse. Cell Rep. Jan 13 2015;10(2):292–305. doi:10.1016/j.celrep.2014.12.014

53. Miura I, Overton ETN, Nakai N, Kawamata T, Sato M, Takumi T. Imaging the Neural Circuit Basis of Social Behavior: Insights from Mouse and Human Studies. Neurol Med Chir (Tokyo). Sep 15 2020;60(9):429–438. doi:10.2176/nmc.ra.2020-0088

54. Vitale EM, Smith AS. Neurobiology of Loneliness, Isolation, and Loss: Integrating Human and Animal Perspectives. Front Behav Neurosci. 2022;16:846315. doi:10.3389/fnbeh.2022.846315

55. Boyko M, Gruenbaum BF, Shelef I, et al. Traumatic brain injury-induced submissive behavior in rats: link to depression and anxiety. Transl Psychiatry. Jun 7 2022;12(1):239. doi:10.1038/s41398-022-01991-1

56. Eyolfson E, Carr T, Khan A, Wright DK, Mychasiuk R, Lohman AW. Repetitive Mild Traumatic Brain Injuries in Mice during Adolescence Cause Sexually Dimorphic Behavioral Deficits and Neuroinflammatory Dynamics. J Neurotrauma. Dec 15 2020;37(24):2718–2732. doi:10.1089/neu.2020.7195

57. Bajwa NM, Halavi S, Hamer M, et al. Mild Concussion, but Not Moderate Traumatic Brain Injury, Is Associated with Long-Term Depression-Like Phenotype in Mice. PLoS One. 2016;11(1):e0146886. doi:10.1371/journal.pone.0146886

58. Wood RL, Thomas RH. Impulsive and episodic disorders of aggressive behaviour following traumatic brain injury. Brain Inj. 2013;27(3):253–61. doi:10.3109/02699052.2012.743181

59. Ozga-Hess JE, Whirtley C, O’Hearn C, Pechacek K, Vonder Haar C. Unilateral parietal brain injury increases risk-taking on a rat gambling task. Exp Neurol. May 2020;327:113217. doi:10.1016/j.expneurol.2020.113217

60. Bhatti JA, Thiruchelvam D, Redelmeier DA. Traumatic brain injury as an independent risk factor for problem gambling: a matched case-control study. Soc Psychiatry Psychiatr Epidemiol. Apr 2019;54(4):517–523. doi:10.1007/s00127-018-1583-1

61. Rostowsky KA, Irimia A, for the Alzheimer’s Disease Neuroimaging I. Acute cognitive impairment after traumatic brain injury predicts the occurrence of brain atrophy patterns similar to those observed in Alzheimer’s disease. GeroScience. 2021/08/01/ 2021;43(4):2015-2039. doi:10.1007/s11357-021-00355-9

62. Madsen T, Erlangsen A, Orlovska S, Mofaddy R, Nordentoft M, Benros ME. Association Between Traumatic Brain Injury and Risk of Suicide. JAMA. Aug 14 2018;320(6):580–588. doi:10.1001/jama.2018.10211

63. Engstrom A, Soderberg S. Transition as experienced by close relatives of people with traumatic brain injury. J Neurosci Nurs. Oct 2011;43(5):253–60. doi:10.1097/JNN.0b013e318227ef9b

64. Gordon WA, Cantor J, Kristen DO, Tsaousides T. Long-term social integration and community support. Handb Clin Neurol. 2015;127:423–31. doi:10.1016/B978-0-444-52892-6.00027-1

65. Silverman JL, Yang M, Lord C, Crawley JN. Behavioural phenotyping assays for mouse models of autism. Nat Rev Neurosci. Jul 2010;11(7):490–502. doi:10.1038/nrn2851

66. Mobbs D, Trimmer PC, Blumstein DT, Dayan P. Foraging for foundations in decision neuroscience: insights from ethology. Nat Rev Neurosci. Jul 2018;19(7):419–427. doi:10.1038/s41583-018-0010-7

67. Leconte C, Benedetto C, Lentini F, et al. Histological and Behavioral Evaluation after Traumatic Brain Injury in Mice: A Ten Months Follow-Up Study. J Neurotrauma. Jun 1 2020;37(11):1342–1357. doi:10.1089/neu.2019.6679

68. Skendelas JP, Muccigrosso M, Eiferman DS, Godbout JP. Chronic Inflammation After TBI and Associated Behavioral Sequelae. Current Physical Medicine and Rehabilitation Report. 2015;3:115–123. 10.1007/s40141-015-0091-4

69. Stoica BA, Faden AI. Cell death mechanisms and modulation in traumatic brain injury. Neurotherapeutics. Jan 2010;7(1):3–12. doi:10.1016/j.nurt.2009.10.023

70. Papadaki E, Kavroulakis E, Manolitsi K, et al. Cerebral perfusion disturbances in chronic mild traumatic brain injury correlate with psychoemotional outcomes. Brain Imaging Behav. Jun 2021;15(3):1438–1449. doi:10.1007/s11682-020-00343-1

71. Kaplan L, Chow BW, Gu C. Neuronal regulation of the blood-brain barrier and neurovascular coupling. Nat Rev Neurosci. Aug 2020;21(8):416–432. doi:10.1038/s41583-020-0322-2

72. Kogan JH, Franklandand PW, Silva AJ. Long-term memory underlying hippocampus-dependent social recognition in mice. Hippocampus. 2000;10(1):47–56. doi:10.1002/(sici)1098-1063(2000)10:1<47::Aid-hipo5>3.0.Co;2-6

73. Tanimizu T, Kenney JW, Okano E, Kadoma K, Frankland PW, Kida S. Functional Connectivity of Multiple Brain Regions Required for the Consolidation of Social Recognition Memory. J Neurosci. Apr 12 2017;37(15):4103–4116. doi:10.1523/JNEUROSCI.3451-16.2017

74. Kim J, Whyte J, Patel S, et al. A perfusion fMRI study of the neural correlates of sustained-attention and working-memory deficits in chronic traumatic brain injury. Neurorehabil Neural Repair. Sep 2012;26(7):870–80. doi:10.1177/1545968311434553

75. Strange BA, Witter MP, Lein ES, Moser EI. Functional organization of the hippocampal longitudinal axis. Nat Rev Neurosci. Oct 2014;15(10):655–69. doi:10.1038/nrn3785

76. de Leon Reyes NS, Sierra Diaz P, Nogueira R, et al. Corticotropin-releasing hormone signaling from prefrontal cortex to lateral septum suppresses interaction with familiar mice. Cell. Sep 14 2023;186(19):4152–4171 e31. doi:10.1016/j.cell.2023.08.010

77. Nolan AL, Sohal VS, Rosi S. Selective Inhibitory Circuit Dysfunction after Chronic Frontal Lobe Contusion. J Neurosci. Jul 6 2022;42(27):5361–5372. doi:10.1523/JNEUROSCI.0097-22.2022

78. Hall ED, Bryant YD, Cho W, Sullivan PG. Evolution of post-traumatic neurodegeneration after controlled cortical impact traumatic brain injury in mice and rats as assessed by the de Olmos silver and fluorojade staining methods. J Neurotrauma. Mar 2008;25(3):235–47. doi:10.1089/neu.2007.0383

79. Saatman KE, Feeko KJ, Pape RL, Raghupathi R. Differential behavioral and histopathological responses to graded cortical impact injury in mice. J Neurotrauma. Aug 2006;23(8):1241–53. doi:10.1089/neu.2006.23.1241

80. Bonne O, Gilboa A, Louzoun Y, et al. Cerebral blood flow in chronic symptomatic mild traumatic brain injury. Psychiatry Res. Nov 30 2003;124(3):141–52. doi:10.1016/s0925-4927(03)00109-4

81. Furuya Y, Hlatky R, Valadka AB, Diaz P, Robertson CS. Comparison of cerebral blood flow in computed tomographic hypodense areas of the brain in head-injured patients. Neurosurgery. Feb 2003;52(2):340–5; discussion 345-6. doi:10.1227/01.neu.0000043931.83041.aa

82. Engel DC, Mies G, Terpolilli NA, et al. Changes of cerebral blood flow during the secondary expansion of a cortical contusion assessed by 14C-iodoantipyrine autoradiography in mice using a non-invasive protocol. J Neurotrauma. Jul 2008;25(7):739–53. doi:10.1089/neu.2007.0480

83. Dixon CE, Kochanek PM, Yan HQ, et al. One-year study of spatial memory performance, brain morphology, and cholinergic markers after moderate controlled cortical impact in rats. J Neurotrauma. Feb 1999;16(2):109–22. doi:10.1089/neu.1999.16.109

84. To XV, Nasrallah FA. A roadmap of brain recovery in a mouse model of concussion: insights from neuroimaging. Acta Neuropathol Commun. Jan 6 2021;9(1):2. doi:10.1186/s40478-020-01098-y

85. Sharp DJ, Graham NSN. Clinical outcomes evolve years after traumatic brain injury. Nat Rev Neurol. Oct 2023;19(10):579–580. doi:10.1038/s41582-023-00868-1

86. Inoue Y, Shiozaki T, Tasaki O, et al. Changes in cerebral blood flow from the acute to the chronic phase of severe head injury. J Neurotrauma. Dec 2005;22(12):1411–8. doi:10.1089/neu.2005.22.1411

87. Hayward NM, Tuunanen PI, Immonen R, Ndode-Ekane XE, Pitkanen A, Grohn O. Magnetic resonance imaging of regional hemodynamic and cerebrovascular recovery after lateral fluid-percussion brain injury in rats. J Cereb Blood Flow Metab. Jan 2011;31(1):166–77. doi:10.1038/jcbfm.2010.67

88. Kochanek PM, Hendrich KS, Dixon CE, Schiding JK, Williams DS, Ho C. Cerebral blood flow at one year after controlled cortical impact in rats: assessment by magnetic resonance imaging. J Neurotrauma. Sep 2002;19(9):1029–37. doi:10.1089/089771502760341947

89. Sharp DJ, Scott G, Leech R. Network dysfunction after traumatic brain injury. Nat Rev Neurol. Mar 2014;10(3):156–66. doi:10.1038/nrneurol.2014.15

90. Woodrow RE, Winzeck S, Luppi AI, et al. Acute thalamic connectivity precedes chronic post-concussive symptoms in mild traumatic brain injury. Brain. Aug 1 2023;146(8):3484–3499. doi:10.1093/brain/awad056

91. Thomas TC, Ogle SB, Rumney BM, May HG, Adelson PD, Lifshitz J. Does time heal all wounds? Experimental diffuse traumatic brain injury results in persisting histopathology in the thalamus. Behav Brain Res. Mar 15 2018;340:137–146. doi:10.1016/j.bbr.2016.12.038

92. Salehi A, Jullienne A, Baghchechi M, et al. Up-regulation of Wnt/beta-catenin expression is accompanied with vascular repair after traumatic brain injury. J Cereb Blood Flow Metab. Feb 2018;38(2):274–289. doi:10.1177/0271678X17744124

93. Golding EM. Sequelae following traumatic brain injury. The cerebrovascular perspective. Brain Res Brain Res Rev. Feb 2002;38(3):377–88. doi:10.1016/s0165-0173(02)00141-8

94. Jullienne A, Obenaus A, Ichkova A, Savona-Baron C, Pearce WJ, Badaut J. Chronic cerebrovascular dysfunction after traumatic brain injury. J Neurosci Res. Jul 2016;94(7):609–22. doi:10.1002/jnr.23732

95. Unterberg AW, Stover J, Kress B, Kiening KL. Edema and brain trauma. Neuroscience. 2004;129(4):1021-9. doi:10.1016/j.neuroscience.2004.06.046

96. Price L, Wilson C, Grant G. Blood-Brain Barrier Pathophysiology following Traumatic Brain Injury. In: Laskowitz D, Grant G, eds. Translational Research in Traumatic Brain Injury. 2016. *Frontiers in Neuroscience*.

97. Zhou Y, Chen Q, Wang Y, et al. Persistent Neurovascular Unit Dysfunction: Pathophysiological Substrate and Trigger for Late-Onset Neurodegeneration After Traumatic Brain Injury. Front Neurosci. 2020;14:581. doi:10.3389/fnins.2020.00581

98. Kenney K, Amyot F, Haber M, et al. Cerebral Vascular Injury in Traumatic Brain Injury. Exp Neurol. Jan 2016;275 Pt 3:353–366. doi:10.1016/j.expneurol.2015.05.019

99. Park E, Bell JD, Siddiq IP, Baker AJ. An analysis of regional microvascular loss and recovery following two grades of fluid percussion trauma: a role for hypoxia-inducible factors in traumatic brain injury. J Cereb Blood Flow Metab. Mar 2009;29(3):575–84. doi:10.1038/jcbfm.2008.151

100. Adams C, Bazzigaluppi P, Beckett TL, et al. Neurogliovascular dysfunction in a model of repeated traumatic brain injury. Theranostics. 2018;8(17):4824–4836. doi:10.7150/thno.24747

101. Bartnik-Olson BL, Holshouser B, Wang H, et al. Impaired neurovascular unit function contributes to persistent symptoms after concussion: a pilot study. J Neurotrauma. Sep 1 2014;31(17):1497–506. doi:10.1089/neu.2013.3213

102. Griffiths DR, Law LM, Young C, et al. Chronic Cognitive and Cerebrovascular Function after Mild Traumatic Brain Injury in Rats. J Neurotrauma. Oct 2022;39(19-20):1429–1441. doi:10.1089/neu.2022.0015

103. Gaggi NL, Ware JB, Dolui S, et al. Temporal dynamics of cerebral blood flow during the first year after moderate-severe traumatic brain injury: A longitudinal perfusion MRI study. Neuroimage Clin. 2023;37:103344. doi:10.1016/j.nicl.2023.103344

104. Dutta S, Sengupta P. Men and mice: Relating their ages. Life Sciences. 2016/05/01/ 2016;152:244-248. 10.1016/j.lfs.2015.10.025

105. Biegon A. Considering Biological Sex in Traumatic Brain Injury. Front Neurol. 2021;12:576366. doi:10.3389/fneur.2021.576366

106. Atlas AR. Mouse Brain [brain atlas]. . atlas.brain-map.org.

107. Tan CO, Meehan WP, 3rd, Iverson GL, Taylor JA. Cerebrovascular regulation, exercise, and mild traumatic brain injury. Neurology. Oct 28 2014;83(18):1665–72. doi:10.1212/WNL.0000000000000944

108. Hamm RJ, Temple MD, O’Dell DM, Pike BR, Lyeth BG. Exposure to environmental complexity promotes recovery of cognitive function after traumatic brain injury. J Neurotrauma. Jan 1996;13(1):41–7. doi:10.1089/neu.1996.13.41

109. Tapias V, Moschonas EH, Bondi CO, et al. Environmental enrichment improves traumatic brain injury-induced behavioral phenotype and associated neurodegenerative process. Exp Neurol. Nov 2022;357:114204. doi:10.1016/j.expneurol.2022.114204

110. Safaryan K, Mehta MR. Enhanced hippocampal theta rhythmicity and emergence of eta oscillation in virtual reality. Nat Neurosci. Aug 2021;24(8):1065–1070. doi:10.1038/s41593-021-00871-z

111. Aida J, Chau B, Dunn J. Immersive virtual reality in traumatic brain injury rehabilitation: A literature review. NeuroRehabilitation. 2018;42(4):441–448. doi:10.3233/NRE-172361

112. Aulisio MC, Han DY, Glueck AC. Virtual reality gaming as a neurorehabilitation tool for brain injuries in adults: A systematic review. Brain Inj. Aug 23 2020;34(10):1322–1330. doi:10.1080/02699052.2020.1802779

113. Ahmadi R, Lim H, Mutlu B, Duff M, Toma C, Turkstra L. Facebook Experiences of Users With Traumatic Brain Injury: A Think-Aloud Study. JMIR Rehabil Assist Technol. Dec 16 2022;9(4):e39984. doi:10.2196/39984

114. Pearce WJ, Doan C, Carreon D, et al. Imatinib attenuates cerebrovascular injury and phenotypic transformation after intracerebral hemorrhage in rats. Am J Physiol Regul Integr Comp Physiol. Dec 1 2016;311(6):R1093–R1104. doi:10.1152/ajpregu.00240.2016

115. Hartmann DA, Coelho-Santos V, Shih AY. Pericyte Control of Blood Flow Across Microvascular Zones in the Central Nervous System. Annu Rev Physiol. Feb 10 2022;84:331–354. doi:10.1146/annurev-physiol-061121-040127

