## Supplemental Material Table and Figures for "Delayed cerebrovascular dysfunction and social deficits after traumatic brain injury"

**Supplementary and Extended Methods**

**Traumatic brain injury (TBI)**

Animals were weighed and placed in a warmed induction chamber for 5minutes at 3% isoflurane (ISOTEC vaporizer). Scalp hair were trimmed, and mice were head fixed via ear bars in a surgical stereotaxic setup (Kopf). Eye ointment (Puralube Vet Ointment, Dechra) was applied, mouse temperature was maintained at 37ºC, and isoflurane at 1.5-2% @ 2L/min during TBI induction (Physiosuite PS0769, Somnosuite, Kent Scientific). Under aseptic conditions, scalp incision was made, and underlying connective tissue was retracted. The craniotomy (center from bregma: AP: –1.25cm, ML: +1.25cm) was etched using an electric dental drill (MH-170, 38000rpm, Foredom), a trephine (5mm diameter), and a drill-bit (0.7mm diameter, #1RF007FST). Cold saline was used intermittently to prevent overheating the skull and underlying tissues. The skull bone (~3mm diameter) was removed, and a 1.5mm impactor tip lowered to exposed brain surface as indicated by the reflective change in pial tissues. The stereotactic arm was adjusted, and the impactor discharged (Leica#39463920, Neuroscience-Tools, 1mm impact-depth, 200ms dwell-time, 5m/s), any extravascular bleeding was wicked away. Incision was closed using either sutures (4mice) or a glass window with dental cement (6mice). Buprenorphine (100ng/g body weight) was injected, and mice were returned to a warmed chamber until ambulatory. Sham mice were anesthetized for a similar duration as the surgery for TBI mice. TBI and sham animals were housed separately (3/cage).

**Behavioral Paradigms**

Animals were handled for ~5 min prior to the onset of each behavioral session. Room ceiling lights were kept on during the behavior (min 250 - max 400 LUX), and fan noise was maintained to minimize distractions. Home cages with mice were held inside the experiment room for 5-10 min before starting the assessments. Mice were allowed to rest for 10min in their home cage between testing paradigms. All setups were wiped clean with cleaning spray (Maxim Germicidal) before and after the session for each animal.

**Behavioral Paradigms**

The behavioral analysis flowchart is shown in Supplementary Fig. 2. Behavior analyses were performed (blinded to injury condition) using Fiji9 (OF, 3Ch) and AnimalTA10 (OF) for animal tracking. Briefly, each video was cropped to isolate the identical regions of interest

(ROIs) for OF and 3Ch. FIJI image adjust algorithm, with automated minimum threshold for animal (red) and background (dark) detection, generated binary masks. Thresholded stacks were averaged as heat-maps (Fiji), with pixel-intensity representing time. Heat-maps allowed estimation of the time spent in each part of the 3Ch arena for assessing relative partner-preference (RPP i.e., the time spent in the partner vs no-partner chamber). Behavior was also scored manually, blind to the test paradigm and conditions using BORIS.11 The absolute partner-interaction time (API i.e. the total time for which the test animal was facing the cylindrical enclosure with the partner) was scored, along with relative partner-interaction time (RPI i.e., the ratio of time spent interacting with partner enclosure vs. total interaction time with both partner and no-partner enclosures).

**MRI Acquisition**

Our perfusion weighted imaging (PWI) MRI methods have been previously published[1]and are as follows. Animals were anesthetized (2% isoflurane with air@ 2L/min, VETQUIP Isoflurane Vaporizer) and their tail vein was cannulated (30gauge) for contrast injection (0.1mmol/kg Gadoterate Meglumine, Dotarem, Guerbet, diluted with sterile saline). Mouse was placed in a custom animal holder (Bruker Biospin) and respiration was monitored with pressure sensor (MP150, BioPAC Systemc Inc.) and maintained at 50-60breaths/minute via isoflurane adjustment (0.25-1%@2L/min, RWD Isoflurane Vaporizer). T2-weighted images (T2WI), T1-weighted images (T1WI) using spin-echo sequence, perfusion-weighted image (PWI) using flow compensated gradient-echo (GEFC) sequence, and susceptibility-weighted images using spine-echo sequence were acquired (Supplementary Table 1). PWI images were acquired with gadolinium injected at 90seconds after sequence onset (volume: 1ul/g of animal weight). Pre-contrast T1WI images were acquired before and post-contrast T1WI images after 10minutes of intravenous injection. The total imaging time was approximately 75minutes. All animals recovered in 5-10minutes after the imaging.

**MRI Image Processing - Perfusion and T2WI**

T2WI were processed using Jim software (V9.1, Xinapse Systems Ltd, UK). T2WI were converted to 'nifti' format using Bruker2Nifti image converter tool.[2] An in-house atlas was prepared using T2WI slices that anatomically resembled the four 1mm thick PWI coronal slices. Specific ROIs based on the mouse Allen brain atlas[3] were manually drawn on selected T2WI using the ROI Analysis tool. For each image set, individual arterial input function (AIF) curves were estimated by automatic AIF selection using the Brain Perfusion tool (# time points in image=150, Time between images=3.947seconds, Arterial and Tissue relaxivity=1.0 L/s/mol, arterial and tissue hematocrit=0.45, SVD threshold=20%). Contrast arrival point was specified using image number. Number of pre-steady-state images and scan number at contrast arrival were estimated through ImageJ based intensity profiles for the PWI image-set. AIFs across all six time points were averaged for each sham animal (Baseline, 3-, 7-, 14-, 30-, 60dpi) to estimate average AIF,[4] further used to estimate the perfusion parameters including cerebral blood flow (CBF-ml/100g-tissue/min) and cerebral blood volume (CBV-%tissue) for each image-set from individual sham animals.[5] Automated AIF was used for TBI-animals to estimate session wise perfusion parameters (CBF, CBV) due to variability in the injury-recovery related factors. Parametric maps were generated for ROI analyses and listed sub regions were delineated: lesion cortex; medial- and lateral-perilesional cortex; regions anterior to lesion: insular visceral and supplementary somatosensory cortex; regions lateral, posterior, and ventral to lesion: adjacent cortex (somatosensory, auditory); distal cortex (rhinal, piriform); and subcortical regions: caudate putamen, dorsal hippocampus, thalamus, and hypothalamus.

**Vessel Painting**

To visualize the cortical angioarchitecture we utilized our vessel painting protocol as previously reported (see Supplementary Materials),[6], Briefly, 1,1'-Dioctadecyl-3,3,3',3'-Tetramethylindocarbocyanine Perchlorate (DiI) was delivered via intracardiac injection prior to , D282, Invitrogen) infusion was performed during transcardial perfusion after the final 60dpi MRI session. Briefly, mice were anesthetized with an intraperitoneal injection of ketamine (90mg/kg) with xylazine (10mg/kg), and analgesia was confirmed with toe pinch. After thoracotomy, DiI solution (0.45mg/ml in phosphate buffered saline (PBS) with 4% dextrose) was manually injected into the left ventricle. Mice were perfused with 10ml PBS and 20 ml 4% paraformaldehyde (PFA) @ 6-7 ml/min using a peristaltic pump (GP1000, Fischer Scientific). After fixation, brains were carefully extracted and post-fixed in 4% PFA for 24hours and stored in PBS until optical imaging. 8/8 TBI and 11/11 sham mice had successful unilateral or bilateral staining of the hemispheres. Vascular topology measurements and analysis were performed by two investigators (one blinded).

**Wide-field fluorescent imaging**

Wide-field cortical images (axial) were acquired with BZ-X810 Keyence microscope (Keyence Corp Osaka, Japan), by positioning the brain between two Superfrost Plus microscope slides

(Fisher Scientific, Pittsburg, PA). High resolution (10x magnification) images of the MCA and its branching vessels from the right hemisphere were acquired.

#### Vascular analysis - Angiotool

ROIs were drawn on the axial cortical images to crop out the lesion (circle) and peri-lesion (torus encapsulating the lesion with same width as the lesion ROI diameter) regions using Fiji software. Angiotool[7] analyses provided vessel density (% area covered by vessels), junctional branch points (branch numbers/mm2), total end points, and average and total length of vessel segments.

#### Fractal analyses

Fractal analyses for vascular complexity was performed using the Fraclac Fiji plugin to obtain the Local fractal dimensions (LFD). Analysis was focused on the lesion and peri-lesion ROIs in 2x axial images[8].

### Supplementary Results

To test the potential of behavior performance at earlier time points in predicting the imminent vascular damage, we measured back-correlations between vascular metrics at 60dpi and social behavior metrics between baseline-30dpi (Fig. 4I, API). In contrast to the 60dpi correlations, the back-correlation shows that sham-vessel density is negatively correlated with API between baseline-7dpi, and positively correlated between 14- and 30dpi. Similar back-correlations were observed for TBI animals during baseline-30dpi. Average vessel length back-correlations with API were positive for sham but negative for TBI animals between baseline-30dpi. Total end point back-correlations for sham animals show positive trend between baseline-7dpi, and negative trend between 14-30dpi, whereas these back-correlations were increasingly positive for TBI animals between baseline-30dpi. Interestingly, maxLFD back-correlations with API were positive for sham and negative for TBI animals. RPI and RPP and back-correlations with vascular metrics show similar trends (Fig. 4I, RPI, RPP).

**Supplementary References**


1. Salehi A, Salari S, Jullienne A, et al. Vascular topology and blood flow are acutely impacted by experimental febrile status epilepticus. *J Cereb Blood Flow Metab*. Jan 2023;43(1):84-98. doi:10.1177/0271678X221117625

2. *Bruker2nifti: Magnetic Resonance Images converter from Bruker ParaVision to Nifti format*. 2017. https://joss.theoj.org/papers/10.21105/joss.00354#

3. Science AIfB. Data from: Allen Mouse Brain Atlas. 2004.

4. Parker GJ, Roberts C, Macdonald A, et al. Experimentally-derived functional form for a population-averaged high-temporal-resolution arterial input function for dynamic contrast-enhanced MRI. *Magn Reson Med*. Nov 2006;56(5):993-1000. doi:10.1002/mrm.21066

5. Petrella JR, Provenzale JM. MR perfusion imaging of the Brain: Techniques and applications. Review. *Am J Roentgenol*. Jul 2000;175(1):207-219. doi:DOI 10.2214/ajr.175.1.1750207

6. Salehi A, Jullienne A, Wendel KM, et al. A Novel Technique for Visualizing and Analyzing the Cerebral Vasculature in Rodents. *Transl Stroke Res*. May 15 2018;doi:10.1007/s12975-018-0632-0

7. Zudaire E, Gambardella L, Kurcz C, Vermeren S. A computational tool for quantitative analysis of vascular networks. *PLoS One*. 2011;6(11):e27385. doi:10.1371/journal.pone.0027385

8. Obenaus A, Ng M, Orantes AM, et al. Traumatic brain injury results in acute rarefication of the vascular network. *Sci Rep*. Mar 22 2017;7(1):239. doi:10.1038/s41598-017-00161-4

9. Schindelin J, Arganda-Carreras I, Frise E, et al. Fiji: an open-source platform for biological-image analysis. *Nat Methods*. Jun 28 2012;9(7):676-82. doi:10.1038/nmeth.2019

10. Chiara V, Kim SY. AnimalTA: A highly flexible and easy-to-use program for tracking and analysing animal movement in different environments. *Methods in Ecology and Evolution*. 2023;14(7):1699-1707. doi:10.1111/2041-210x.14115

11. Friard O, Gamba M, Fitzjohn R. BORIS: a free, versatile open-source event-logging software for video/audio coding and live observations. *Methods in Ecology and Evolution*. 2016;7(11):1325-1330. doi:10.1111/2041-210x.12584

**Supplementary Figures**

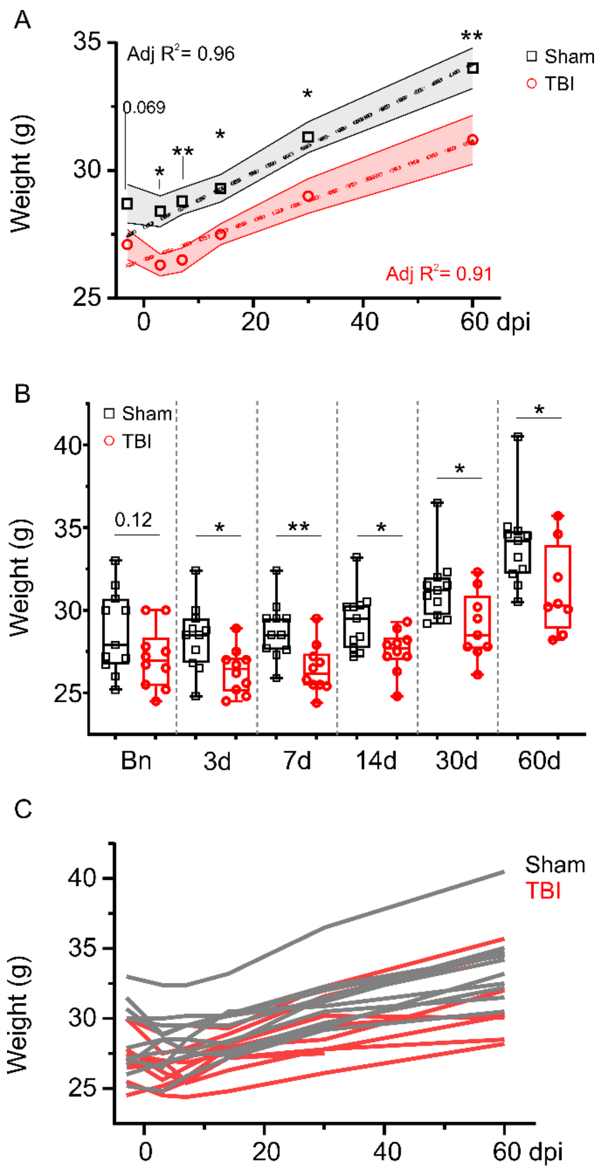


**Supplemental Figure 1. Animal weights trend during the time course**. **A)** Average weights for sham (black, square) and TBI mice (red, circle), shaded area – SEM (2w ANOVA, Injury F(1,111)=35.5, ***p<0.0001; timepoint F(5,111)=19.8, ***p<0.0001. Interaction – ns. Tukey's Post hoc: baseline – trending, p=0.069; 3dpi – *p=0.018; 7dpi – **p=0.009; 14dpi – *p=0.04; 30dpi – *p=0.0115; 60dpi – **p=0.003). Dashed lines represent linear fit with similar increases in body weight. Sham average weight profile shows relatively higher weights, including baseline, likely due to the outliers as shown in panel C. **B)** Boxplots of time point comparisons between sham and TBI mice (1w ANOVA for longitudinal comparison, no significant differences; 2-tailed unpaired t-tests for each time point comparison: Baseline – t=1.62, df=19, p=0.12; 3dpi – t=2.72, df=19, *p=0.0135; 7dpi – t=3.30, df=19, **p=0.004; 14dpi – t=2.64, df=19, *p=0.016; 30dpi – t=2.54, df=18, *p=0.02; 60dpi – t=2.22, df=17, *p=0.04). **C)** Spaghetti plots show trends for individual animals.

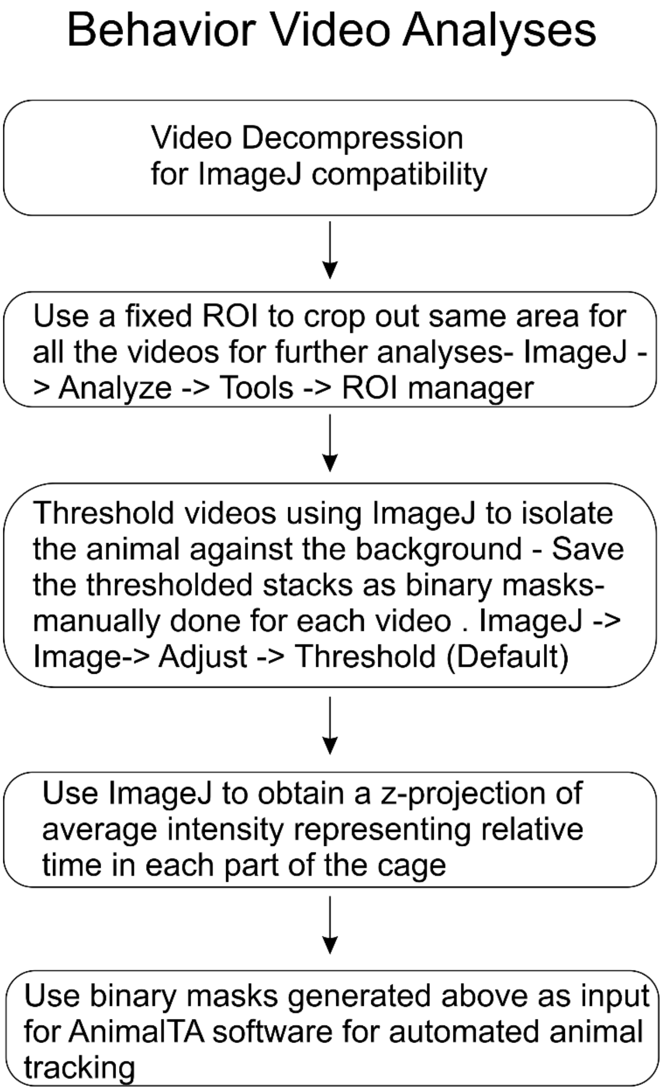


**Supplementary Figure 2.** Flow chart of steps for semi-automated image analysis for open field and 3Ch behavior videos.

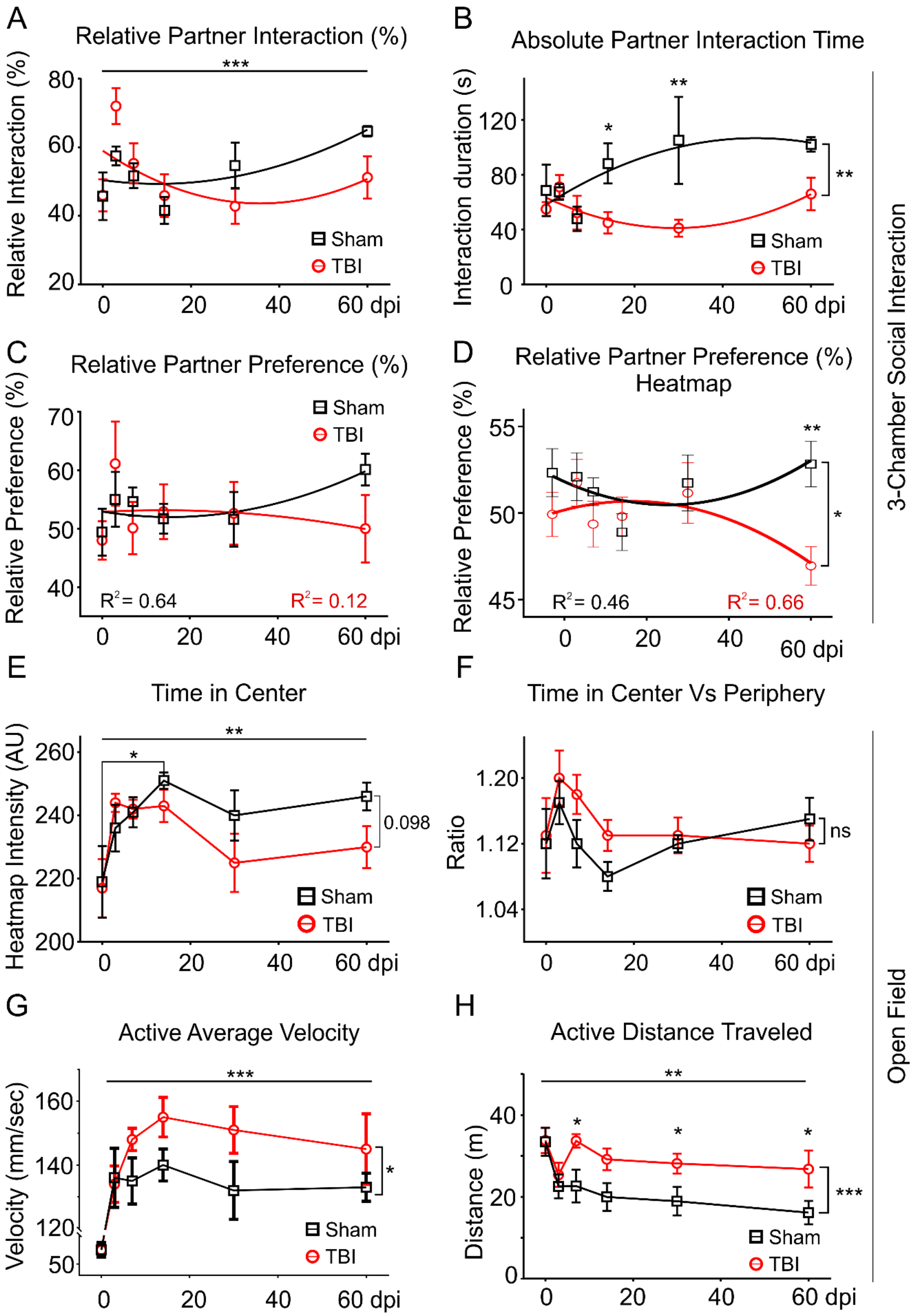

A
Relative Partner Interaction (%)
Relative Interaction (%)
***
Sham
TBI
0 20 40 60 dpi
B
Absolute Partner Interaction Time
Interaction duration (s)
*
**
**
Sham
TBI
0 20 40 60 dpi
C
Relative Partner Preference (%)
Relative Preference (%)
Sham
TBI
R² = 0.64
R² = 0.12
0 20 40 60 dpi
D
Relative Partner Preference (%) Heatmap
Relative Preference (%)
**
*
R² = 0.46
R² = 0.66
0 20 40 60 dpi
3-Chamber Social Interaction
E
Time in Center
Heatmap Intensity (AU)
**
*
0.098
Sham
TBI
0 20 40 60 dpi
F
Time in Center Vs Periphery
Ratio
ns
Sham
TBI
0 20 40 60 dpi
G
Active Average Velocity
Velocity (mm/sec)
***
*
Sham
TBI
0 20 40 60 dpi
H
Active Distance Traveled
Distance (m)
**
*
*
*
***
Sham
TBI
0 20 40 60 dpi
Open Field

**Supplemental Figure 3. Additional metrics from open field (A-D) and 3-chamber social interaction behavior (E-H).** **A)** Relative partner interaction (ratio of time interacting with partner vs total interaction time interacting with both the cylindrical enclosures (occupied and empty) was significantly different across timepoints (F (5, 61) = 4.78, p = 0.0009) but similar across injury condition (p = 0.88). Post-hoc, differences across timepoints were driven for shams by higher relative partner interaction at 60dpi vs 14dpi (*p = 0.03). For TBI animals, the differences across time are driven by higher partner interaction at 3dpi vs baseline (**p = 0.009), 14dpi (**p = 0.008), and 30dpi (**p = 0.002). **B)** Absolute interaction time was higher for sham compared to TBI animals (injury factor only, (F (1, 60) = 10.0, p = 0.002), post hoc driven by higher sham values at 14 dpi (*p = 0.028) and 30 dpi (**p = 0.001). **C)** Relative partner preference (RPP) based on the time spent in the partner's chamber (manual scoring) show a non-significant delayed decline in RPP for TBI mice compared to sham animals (ns). **D)** Semi-automated estimation of RPP based on heatmap analyses found a significant differences between sham and TBI mice (F (1, 60) = 5.02, *p = 0.029), post-hoc driven by TBI dependent reduced preference for partner at 60 dpi as compared to shams (**p = 0.002). **E)** Automated animal tracking revealed significant decreases in time spent by TBI (red) and sham (black) mice in center of the open field arena across timepoints (F (5, 62) = 4.50, p = 0.001) but similar across injury condition (sham vs TBI, F (1, 62) = 1.92, p = 0.17). Post-hoc comparison found no significant differences in center-zone exploration across timepoints for both sham and TBI mice with trending decrease at 60 dpi (p = 0.098) in TBI mice. **F)** The ratio of time spent in center vs periphery for sham and TBI mice was not significantly different across timepoints or injury conditions. **G)** Active average velocity while animals were moving was significantly different across timepoints (F (5, 61) = 41.50, p < 0.001) and injury condition (F (1, 61) = 5.36, p = 0.024), post-hoc driven only due the difference between baseline and 3dpi for both sham and TBI mice. For each timepoint, baseline and TBI values were similar. Baseline was the first exposure to behavior set up and animals were cautious due to neophobia. Low baseline average active velocity indicates shorter bouts of movements compared to other sessions where animals show longer bouts after gaining familiarity with the set up. **H)** Distance moved by TBI animals during active exploration was higher compared to shams (F (1, 61) = 15.1, p = 0.0003), across several timepoints (F (5, 61) = 3.67, p = 0.006). Post-hoc, the differences were driven by higher values for TBI mice compared to shams at 7- (*p = .015), 14- (*p = 0.049), 30-(*p = 0.048), and 60 dpi (*p = 0.023).

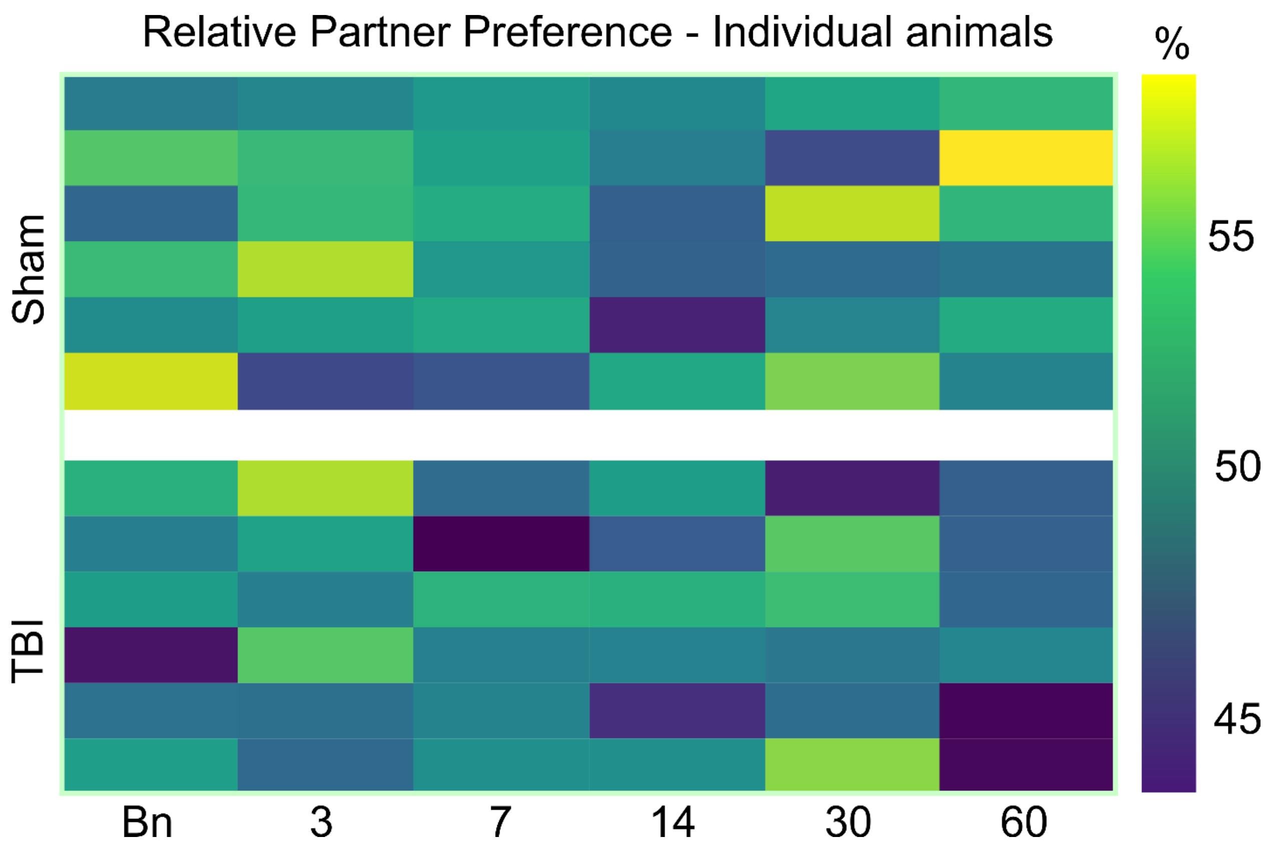


**Supplemental Figure 4.** Heatmap representing the relative partner preference (semi-automated) for each sham and TBI animal across all time points with a decline in RPP at 60dpi, injured mice.

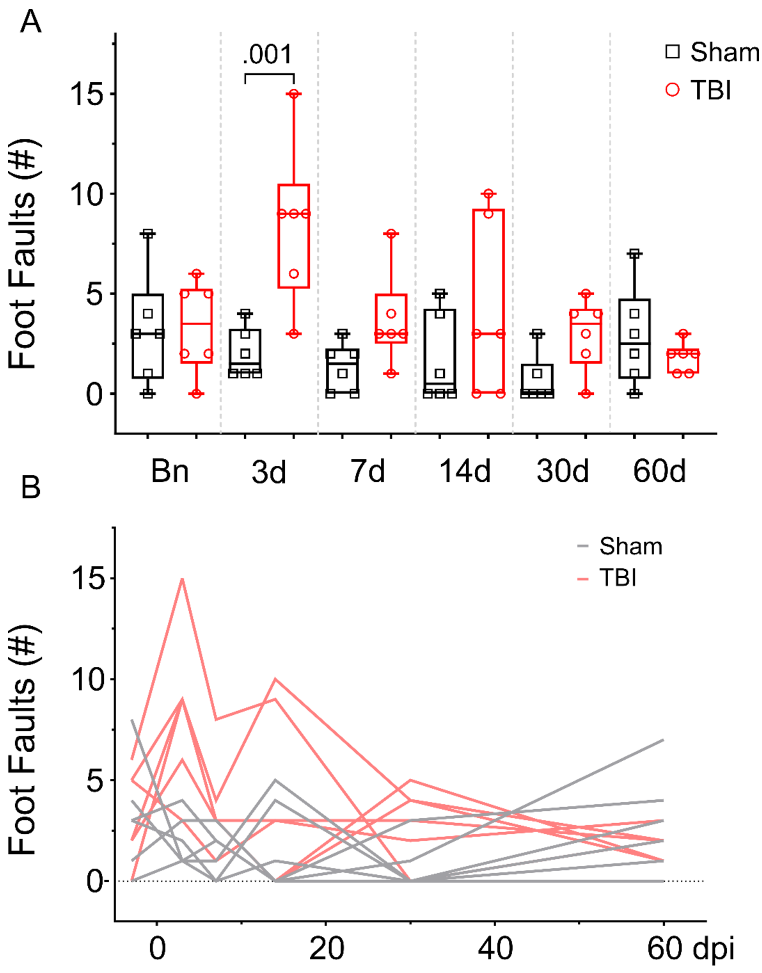


**Supplemental Figure 5** – **A)** Foot fault behavior in sham and TBI mice across time. TBI mice exhibited higher motor impairment at 3dpi with gradual recovery over time. 1wANOVA, $F(11, 60) = 3.99$, ***$p < 0.001$, Tukey's post hoc 3dpi sham vs TBI **$p = 0.001$. **B)** Spaghetti plots showing the foot fault trends for individual animals.

**Supplementary Table 1: 9.4T Neuroimaging parameters**

| MRI Sequence | TE (ms) | TR (ms) | NEX | FOV ($mm^2$) | Dim. | Slices | Slice Thick-ness (mm) | In-plane Resolution ($mm^2$) | Imaging time (min) |
|---|---|---|---|---|---|---|---|---|---|
| T2WI | 10 | 4000 | 4 | 15x15 | $128^2$ | 30 | 0.5 | $0.117^2$ | 25.6 |
| T1WI | 5.2 | 22.3 | 1 | 15x15 | $128^2$ | 64 | 0.25 | $0.117^2$ | 5.0 |
| GEFC-PWI | 3.5 | 61.7 | 1 | 15x15 | $64^2$ | 4 | 1.0 | $0.234^2$ | 9.8 |
| SWI | 10.7 | 723 | 8 | 15x15 | $128^2$ | 30 | 0.5 | $0.117^2$ | 12.6 |

Dim. – acquisition dimensionality (pixels), FOV – field of view, NEX – number of averages, GEFC-PWI – Flow compensated gradient echo-perfusion weighted image, SWI – Susceptibility weighted image, T1WI – T1-weighted image, T2WI – T2-weighted image, TE – Time to echo, TR – Repetition time,
